# Conformational and dynamical plasticity in substrate-binding proteins underlies selective transport in ABC importers

**DOI:** 10.1101/537308

**Authors:** Marijn de Boer, Giorgos Gouridis, Ruslan Vietrov, Stephanie L. Begg, Gea K. Schuurman-Wolters, Florence Husada, Nikolaos Eleftheriadis, Bert Poolman, Christopher A. McDevitt, Thorben Cordes

**Affiliations:** Molecular Microscopy Research Group, Zernike Institute for Advanced Materials, University of Groningen, Nijenborgh 4, 9747 AG Groningen, The Netherlands; Physical and Synthetic Biology, Faculty of Biology, Ludwig Maximilians-Universität München, Großhadernerstr. 2-4, 82152 Planegg-Martinsried, Germany; KU Leuven, Department of Microbiology and Immunology, Rega Institute for Medical Research, Laboratory of Molecular Bacteriology, 3000 Leuven, Belgium; Department of Biochemistry, Groningen Biomolecular Science and Biotechnology Institute & Zernike Institute for Advanced Materials, University of Groningen, Nijenborgh 4, 9747 AG Groningen, The Netherlands; Department of Microbiology and Immunology, The Peter Doherty Institute for Infection and Immunity, University of Melbourne, Melbourne, Victoria, 3000, Australia; Research Centre for Infectious Diseases, School of Biological Sciences, The University of Adelaide, Adelaide, South Australia, 5005, Australia

**Keywords:** ABC transporters, conformational dynamics, single-molecule FRET, substrate-binding proteins, solute translocation, substrate transport

## Abstract

Substrate-binding proteins (SBPs) are associated with ATP-binding cassette importers and switch from an open-to a closed-conformation upon substrate binding providing specificity for transport. We investigated the effect of substrates on the conformational dynamics of six SBPs and the impact on transport. Using single-molecule FRET, we reveal an unrecognized diversity of plasticity in SBPs. We show that a unique closed SBP conformation does not exist for transported substrates. Instead, SBPs sample a range of conformations that activate transport. Certain non-transported ligands leave the structure largely unaltered or trigger a conformation distinct from that of transported substrates. Intriguingly, in some cases similar SBP conformations are formed by both transported and non-transported ligands. In this case, the inability for transport arises from slow opening of the SBP or the selectivity provided by the translocator. Our results reveal the complex interplay between ligand-SBP interactions, SBP conformational dynamics and substrate transport.

## INTRODUCTION

ATP-binding cassette (ABC) transporters facilitate the unidirectional trans-bilayer movement of a diverse array of molecules using the energy released from ATP hydrolysis^1^. ABC transporters share a common architecture, with the translocator unit comprising two transmembrane domains (TMDs) that form the translocation pathway and two cytoplasmic nucleotide-binding domains (NBDs) that bind and hydrolyse ATP. ABC importers require an additional extra-cytoplasmic accessory protein referred to as a substrate-binding protein SBP or domain SBD (hereafter SBDs and SBPs are both termed SBPs)^2–4^. ABC importers that employ SBPs can be subdivided as Type I or Type II based on structural and mechanistic distinctions^5, 6^. A unifying feature of the transport mechanism of Type I and Type II ABC importers is the binding and delivery of substrate from a dedicated SBP to the translocator unit for import into the cytoplasm.

Bacterial genomes encode multiple distinct ABC importers to facilitate the acquisition of essential nutrients such as sugars, amino acids, vitamins, compatible solutes, and metal ions^1, 7^. Many ABC importers can transport more than one substrate using high-affinity interactions between SBPs and transported ligands (herein termed cognate substrates)^2^. Despite low sequence similarity between SBPs of different ABC importers, they share a common architecture comprising two structurally conserved rigid lobes connected by a flexible hinge region (**Figure 1**)^2^. Numerous biophysical^8^ and structural analyses^9^ indicate that ligand binding at the interface of the two lobes facilitates switching between two conformations, i.e. from an open to a closed conformation. Bending and unbending of the hinge region brings the two lobes together (closed conformation) or apart (open conformation), respectively. Crystallographic analysis show that the amount of opening varies between different SBPs; the lobe-movements observed range from small rearrangements as in the Type II SBP BtuF^10^, to complete reorientation of both lobes by angles as large as 60° in the Type I SBP LivJ^11^. Nevertheless, the wealth of structural data permits a structural classification of SBPs, wherein the hinge region is the most defining feature of each sub-group or cluster (**Figure 1**)^2, 3^. Crystal structures of the same protein, but with different ligands bound, generally report the same degree of closing of the SBP^11–15^.

**Figure 1.**
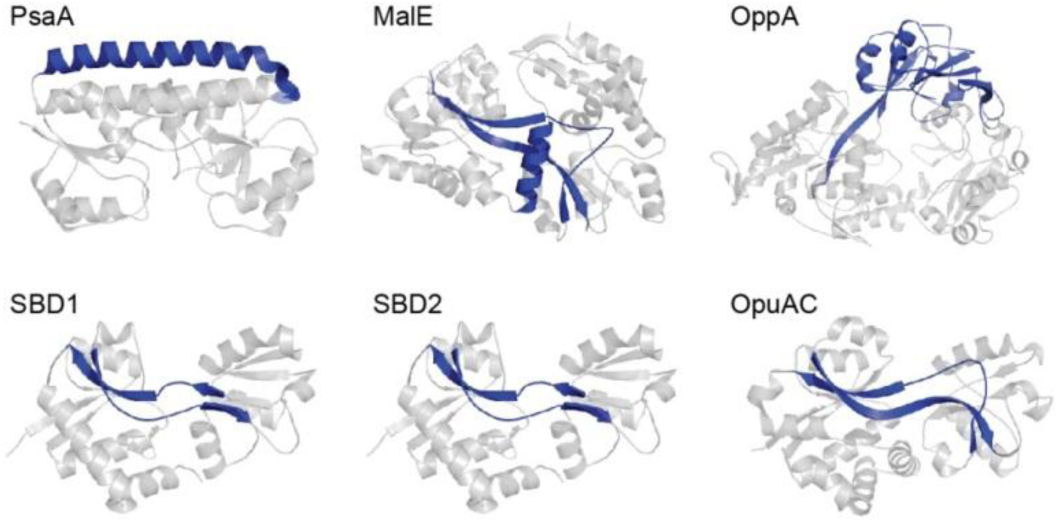
Representative SBPs from different structural clusters, categorized by their hinge region. X-ray crystal structures of PsaA (3ZK7; cluster A), MalE (1OMP; cluster B), OppA (3FTO; cluster C), OpuAC (3L6G; cluster F), SBD1 (4LA9; cluster F) and SBD2 (4KR5; cluster F) are all shown in the open ligand-free conformation. Hinge regions are shown in blue and the two rigid lobes in grey. For classification of the proteins in clusters see Berntsson *et al* and Scheepers *et al*^2, 3^.

It thus is assumed that the conformational switching of the SBPs enables the ABC transporter to allosterically sense the loading state of the SBP-ligand complex (‘translocation competency’), thereby contributing to transport specificity^7,9^. For example, crystal structures of the SBP MalE show that the protein adopts a unique closed conformation when interacting with cognate ligands maltose, maltotriose and maltotetraose^15^, while the non-transported ligand β-cyclodextrin is bound by MalE^16,17^but fails to trigger formation of the closed conformation^17–19^. Ligands that are bound by the SBP, but not transported, are termed herein non-cognate ligands. Such findings suggest that only SBPs that adopt the closed conformation can productively interact with the translocator and initiate transport. However, the TMDs of certain ABC importers were also shown to interact directly with their substrates. In MalFGK_2_E^20^ from *Escherichia coli* and Art(QM)_2_^21^ from *Thermoanaerobacter tengcongensi* substrate-binding pockets have been identified inside the TMDs, and these might be linked to regulation of transport. Similar binding pockets within the TMDs have not been observed in the high-resolution structures of other ABC importers, although cavities through which the substrate passes in the transition of the TMD from outward-to inward-facing must be present in all the transporters^22–24^. Additional complexity exists for the coupling of SBP conformational switching and the ligand recognition process, as crystallographic^25, 26^, nuclear magnetic resonance (NMR)^27^ and single-molecule^28, 29^ studies indicate that SBPs can undergo intrinsic conformational changes in the absence of substrate. Furthermore, crystal structures of the SBPs MalE and a D-xylose SBP in an open ligand-bound conformation were obtained^30, 31^. Such observations question the precise relationship between SBP-ligand interactions, SBP conformational changes and their involvement in transport function.

A range of biophysical and structural approaches have already been used to decipher the mechanistic basis of SBP-ligand interactions^8, 9, 11, 17, 19^. However, these techniques only provide information on the overall population of molecules. Recent advances in single-molecule methodologies now permit new insight into the conformational heterogeneity, dynamics and occurrences of rare events in SBPs^28, 29, 32–35^, which are difficult to obtain in bulk measurements. Here, we combined single-molecule Förster resonance energy transfer (smFRET)^36^ and transport measurements to investigate how cognate and non-cognate substrates influence the conformational states and the underlying dynamics of SBPs. Six distinct SBPs were selected (**Figure 1**)^37–41^, based on two criteria. First, they cover the breadth of SBP structural classes: PsaA (cluster A), MalE (cluster B), OppA (cluster C), SBD1 and SBD2 of GlnPQ, and OpuAC (all cluster F). The selected SBPs provide coverage of hinge region diversity^2, 3^, thereby addressing a hypothesized key determinant in SBP conformational dynamics. Moreover, subtle structural or sequence differences among SBPs that belong to the same cluster are addressed by examining SBD1, SBD2 and OpuAC that all belong to cluster F. Second, the selected SBPs belong to Type I and Type II ABC importers with extensively characterized substrate (cognate and non-cognate) interactions, such as metal ions (PsaA)^40^, sugars (MalE)^42^, peptides (OppA)^43^, amino acids (SBD1 and SBD2)^37^, and compatible solutes (OpuAC)^38^.

## RESULTS

### Multiple SBP conformations are translocation competent

Crystal structures of SBPs suggest that ligand binding is coupled to switching between two protein conformations, an open and a closed conformation. Mechanistically this process has been linked to the allosteric regulation of substrate transport^7–9, 44–48^. Here, we assessed this model by investigating the interaction of six SBPs, PsaA, MalE, OppA, SBD1, SBD2 and OpuAC, with a range of cognate substrates. We employed single-molecule FRET to analyse SBP conformations, wherein each of the two SBP lobes was labelled with either a donor or an acceptor fluorophore (**Figure 2A**)^29, 49^. Surface-exposed and non-conserved residues, showing largest distance changes according to the crystal structures of the open and closed states, were chosen as suitable cysteine positions for labelling. Protein labelling did not alter the ligand-binding affinity, that is, the ligand dissociation constant K_D_ (**Table 1**). In our assays, the inter-dye separation reports on the relative orientation and distance between the SBP lobes and is thus indicative for the degree of closing. Steady-state anisotropy measurements indicate that the dyes retain sufficient rotational freedom (**Table 2**) so that relative inter-dye distances can be accessed via the apparent FRET efficiency of freely diffusing or surface-immobilized protein molecules. Although this approach monitors only a single distance in the SBP, it permits rapid screening of ligand induced conformational changes in physiologically relevant conditions.

**Table 1.** Dissociation constants of substrate-binding proteins.

| Protein<sup>d</sup> | Ligand | K<sub>D</sub> labelled protein<sup>e</sup> (μM) | K<sub>D</sub> WT protein<sup>f</sup> (μM) |
| --- | --- | --- | --- |
| OpuAC(V360C/N423C) | Glycine betaine | 3.4 ± 0.4<sup>a</sup> | 4-5<sup>38</sup> |
| OppA(A209C/S441C) | RPPGFSFR | 14 ± 3<sup>b</sup> | 5 ± 3<sup>b</sup> |
| SBD2(T369C/S451) | Glutamine | 1.2 ± 0.2<sup>c</sup> | 0.9 ± 0.1<sup>29</sup> |
| SBD1(T159C/G87C) | Asparagine | 0.34 ± 0.03<sup>c</sup> | 0.2 ± 0.0<sup>29</sup> |
| MalE(T36C/S352C) | Maltose | 1.7 ± 0.3<sup>a</sup> | 1<sup>42</sup> |

**Table 2.** Steady-state anisotropy values.

|  | Anisotropy |  |  |  |
| --- | --- | --- | --- | --- |
|  | Alex555 | Alexa647 | Cy3B | Atto647N |
| Free dye | 0.25 | 0.20 | 0.08 | 0.08 |
| OpuAC(V360C/N423C) | NA | NA | 0.17 | 0.11 |
| OppA(A209C/S441C) | 0.25 | 0.19 | NA | NA |
| SBD1(G87C/T159C) | 0.27 | 0.19 | NA | NA |
| SBD2(T369C/S451) | 0.26 | 0.20 | NA | NA |
| MalE(T36C/S352C) | 0.29 | 0.24 | NA | NA |
| PsaA(V76C/K237C) | 0.28 | 0.22 | NA | NA |

NA: not applicable. Data correspondents to mean (s.d. below <0.01) of duplicate experiments with the same labelled protein sample.

**Figure 2.**
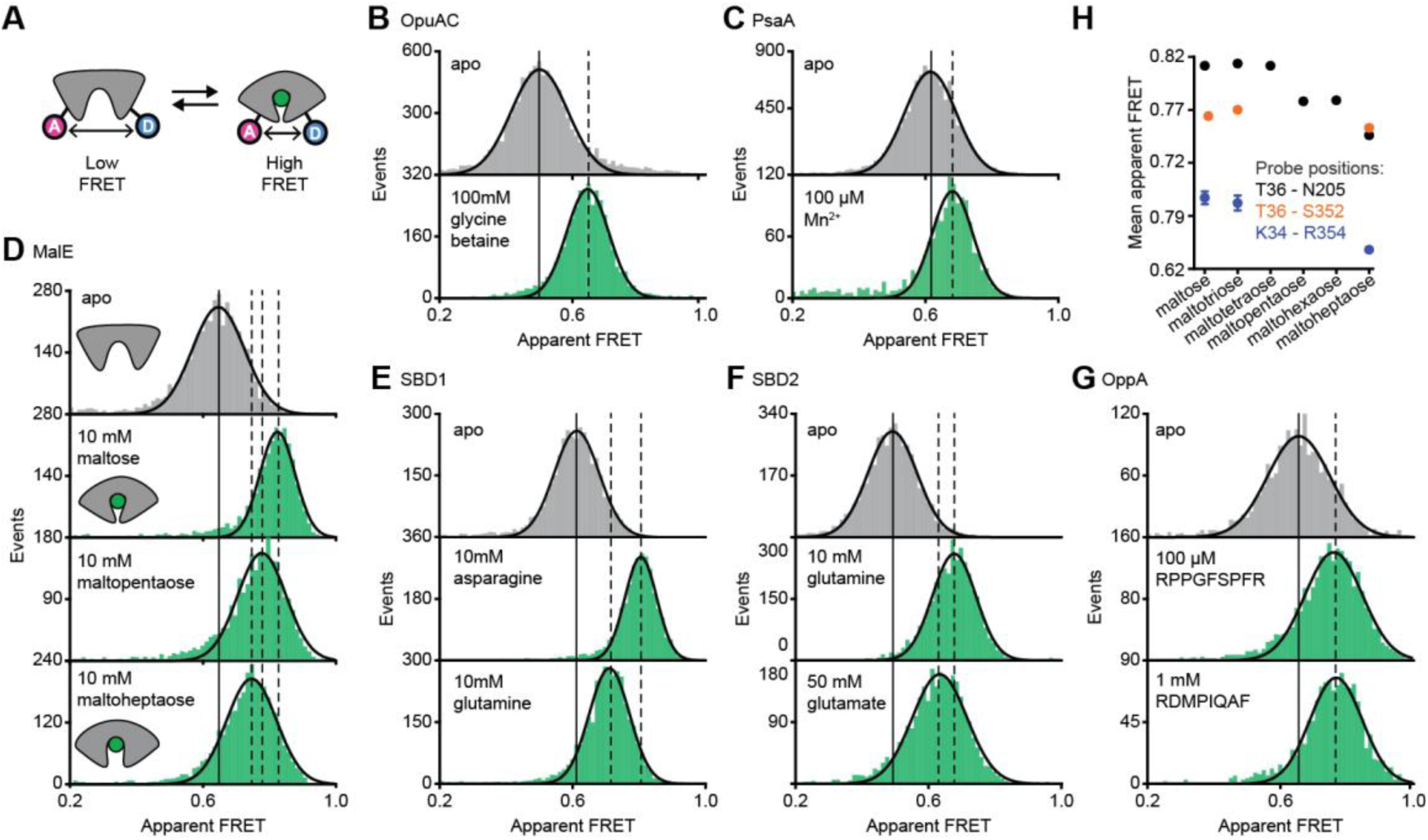
Conformational states of SBPs probed by smFRET reveal multiple active conformations. (**A**) Experimental strategy to study SBP conformational changes via FRET. Solution-based apparent FRET efficiency histograms of OpuAC(V360C/N423C) (**B**), PsaA(V76C/K237C) (**C**), MalE(T36C/S352C) (**D**), SBD1(T159C/G87C) (**E**), SBD2(T369C/S451) (**F**) and OppA(A209C/S441C) (**G**) in the absence (grey bars) and presence of different cognate substrates (green bars). The OppA substrates are indicated by one-letter amino acid code. Bars are experimental data and the solid line a Gaussian distribution fit. The 95% confidence interval of the Gaussian distribution mean is shown in Table 3, and the interval centre is indicated by vertical lines (solid and dashed). (**H**) Mean of the Gaussian distribution of MalE labelled at T36/S352 (black), T36/N205 (green) or K34/R352 (orange). Error bars indicate 95% confidence interval of the mean.

The apparent FRET efficiency of individual and freely-diffusing SBPs were measured in the presence and absence of their cognate substrates by using confocal microscopy. Saturating concentrations of cognate substrate, above the dissociation constant K_D_ (**Table 1**), shift the FRET efficiency histograms and the fitted Gaussian distributions to higher values compared to the ligand-free SBPs (**Figure 2B-G**; **Table 3**), indicating a reduced distance between the SBP lobes and thus closure of the proteins. The solution-based FRET distributions of ligand-bound and ligand-free SBPs are unimodal and thus do not reveal any substantial conformational heterogeneity, such as a pronounced closing in the absence of substrate or a substantial population of an open-liganded state (*vide infra*). This strongly suggests that ligands are bound via an induced-fit mechanism, unless dynamics occurs on timescales faster than milliseconds. This inference was confirmed for OppA by examining individual surface-immobilized proteins and demonstrating that substrate-induced SBP closing follows first-order kinetics while the opening obeys zeroth-order kinetics (**Figure 2 – figure supplements 1**)^32^.

**Table 3.** Apparent FRET efficiency values of solution-based measurements.

| Protein | Condition | Apparent FRET* |
| --- | --- | --- |
| OpuAC(V360C/N423C) | no ligand | 0.501 ± 0.004 |
|  | proline | 0.479 ± 0.003 |
|  | carnitine | 0.543 ± 0.004 |
|  | glycine betaine | 0.646 ± 0.003 |
| OppA(A209C/S441C) | no ligand | 0.655 ± 0.005 |
|  | RPPGFSPFR | 0.767 ± 0.004 |
|  | RDMP IQAF | 0.760 ± 0.004 |
|  | SLSQSKVLPVPQ | 0.761 ± 0.006 |
|  | SLSQSKVLP | 0.770 ± 0.009 |
| SBD2(T369C/S451) | no ligand | 0.492 ± 0.003 |
|  | glutamate | 0.633 ± 0.004 |
|  | glutamine | 0.677 ± 0.002 |
|  | arginine | 0.496 ± 0.002 |
| SBD1(G87C/T159C) | no ligand | 0.612 ± 0.003 |
|  | glutamine | 0.710 ± 0.003 |
|  | asparagine | 0.805 ± 0.002 |
|  | histidine | 0.761 ± 0.003 |
|  | arginine | 0.610 ± 0.002 |
|  | lysine | 0.613 ± 0.002 |
| MalE(T36C/S352C) | no ligand | 0.646 ± 0.003 |
|  | B-cyclodextrin | 0.696 ± 0.003 |
|  | maltotriitol | 0.725 ± 0.003 |
|  | maltotetraitol | 0.725 ± 0.003 |
|  | maltose | 0.824 ± 0.002 |
|  | maltotriose | 0.826 ± 0.003 |
|  | maltotetraose | 0.824 ± 0.002 |
|  | maltopentaose | 0.777 ± 0.004 |
|  | maltohexaose | 0.779 ± 0.003 |
|  | maltoheptaose | 0.746 ± 0.002 |
|  | maltooctaose | 0.745 ± 0.003 |
|  | maltodecaose | 0.742 ± 0.003 |
| MalE(T36C/N205C) | no ligand | 0.538 ± 0.007 |
|  | maltoheptaose | 0.755 ± 0.005 |
|  | maltooctaose | 0.758 ± 0.005 |
|  | maltodecaose | 0.757 ± 0.006 |
| MalE(K34C/R354C) | no ligand | 0.524 ± 0.010 |
|  | maltoheptaose | 0.669 ± 0.008 |
|  | maltooctaose | 0.662 ± 0.007 |
|  | maltodecaose | 0.666 ± 0.008 |
| PsaA(V76C/K237C) | no ligand | 0.615 ± 0.003 |
|  | Mn<sup>2+</sup> | 0.681 ± 0.004 |
|  | Zn<sup>2+</sup> | 0.687 ± 0.005 |
| PsaA(E74C/K237C) | no ligand | $0.518 \pm 0.003$ |
| | Mn<sup>2+</sup> | $0.567 \pm 0.003$ |
| | Zn<sup>2+</sup> | $0.570 \pm 0.003$ |
| PsaA(D280N/V76C/K237C) | no ligand | $0.617 \pm 0.003$ |
| | Zn<sup>2+</sup> | $0.688 \pm 0.006$ |

\*95% confidence interval for the mean of the Gaussian distribution fit
\*\*Only data of the same protein construct can be compared (all recorded on one day), due to unavoidable differences in microscope settings when measurements are done on different days. Qualitatively consistent results were obtained upon repetition with an independent labelled protein sample and measured on a different day.

Further examination of the FRET distributions shows that multiple substrate-bound SBP conformations exist for SBD1, SBD2 and MalE (**Figure 2D-F**). For the amino acid binding-proteins SBD1 and SBD2, the cognate substrates^37^ asparagine and glutamine for SBD1, and glutamine and glutamate for SBD2 all stabilize a distinct protein conformation, as the FRET efficiency histograms and the fitted Gaussian distributions are different (**Figure 2E-F**; **Table 3**). For the maltodextrin binding-protein MalE we examined the effect of cognate maltodextrins^39^, ranging from two to seven glucosyl units, on the MalE conformation. Comparison of the FRET efficiency histograms of the different MalE-ligand complexes shows that at least three distinct ligand-bound MalE conformations exist (**Figure 2D**; **Figure 2 – figure supplements 2A**). Contrary to SBD1 and SBD2, some cognate substrates did not induce a unique MalE conformation. For example, maltopentaose and maltohexaose elicited the same FRET change, and triggered the formation of a partial closed MalE conformation (**Figure 2 – figure supplements 2A**). However, this conformational state is different from the full closed form of MalE, which is obtained by maltose, maltotriose and maltotetraose, or the other partial closed conformation that is formed by binding of maltoheptaose (**Figure 2 – figure supplements 2A**). The results for MalE were confirmed by examining different inter-dye positions (**Figure 2H**; **Figure 2 – figure supplements 2**).

However, whether this conformational plasticity is a universal feature among SBPs needs to be tested further, because in OppA the four examined cognate substrates^43^ elicited the same FRET change (**Figure 2G**; **Figure 2 – figure supplements 2B**). Taken together, these data indicate that although the examined SBPs have a single open conformation, a productive interaction between the SBP and the translocator does not require a single, unique closed SBP conformation. The structural flexibility of the SBP permits the formation of one or more ligand-bound conformations, all of which are able to interact with the translocator and initiate transport^37-40, 43^.

### Intrinsic conformational changes of SBPs

We then investigated whether the conformational changes in the SBPs that were triggered by their ligands, can also occur in their absence. To address this, we investigated surface-tethered SBPs in the absence of ligand and used confocal scanning microscopy to obtain millisecond temporal resolution. Compared to the solution-based smFRET experiments, individual surface-tethered SBPs greatly increase the sensitivity to detect rare events. In contrast to prior work^28, 29, 32, 33^, the labelled SBPs were supplemented with high concentrations of unlabelled protein (20 μM), or the divalent chelating compound ethylenediaminetetraacetic acid (EDTA, [c] = 1 mM for PsaA), to remove any contaminating ligands (**Figure 3A**). Contaminations could otherwise lead to conformational changes that are misinterpreted as intrinsic closing of the SBP. Consistent with the solution-based measurements, all SBPs were predominately in a low FRET state (open conformation; **Figure 3B-G**; **Figure 3 – figure supplements 3**). For ligand-free MalE, PsaA and OpuAC, no transitions to higher FRET states were observed within a total observation time of >8 min for each SBP (**Figure 3B-D**; **Table 4**). In SBD1, SBD2 and OppA rare transitions to a high FRET state can be observed and have an average lifetime of 110 ± 14, 77 ± 7 and 230 ± 50 ms (mean ± s.e.m.) for SBD1, SBD2 and OppA, respectively (**Figure 3E-G**; **Figure 3 – figure supplements 3D-F**). Transitions towards these states occur only rarely, i.e. on average every 15, 10 or 20 s for SBD1, SBD2 and OppA, respectively (**Figure 3H**; **Table 4**). Taken together, some SBPs have the ability to also close without the ligand on the second timescale. However, not all SBPs show intrinsic conformational transitions, unless these occur below the temporal resolution of the measurements (millisecond timescale). Overall, the data indicate that diversity exists in the conformational dynamics of ligand-free SBPs.

**Table 4.** Statistics of confocal scanning experiments of immobilized molecules.

| Protein* | Condition | Molecules analysed | Total observation time (min) | Transitions observed |
| --- | --- | --- | --- | --- |
| OpuAC(V360C/N423C) | no ligand | 702 | 8.2 | 0 |
|  | 1 mM Glycine Betaine | 78 | 2.3 | 0 |
| OppA(A209C/S441C) | no ligand | 326 | 11.0 | 37 |
| | 2.5 μM RPPGFSPFR | 39 | 1.4 | 329 |
| | 5 μM RPPGFSPFR | 168 | 6.0 | 2166 |
| | 10 μM RPPGFSPFR | 29 | 0.7 | 280 |
| | 20 μM RPPGFSPFR | 65 | 2.4 | 1837 |
| | 200 μM RPPGFSPFR | 31 | 0.5 | 0 |
| SBD1(G87C/T159C) | no ligand | 384 | 16.1 | 67 |
|  | 1 mM asparagine | 72 | 8.0 | 0 |
| SBD2(T369C/S451) | no ligand | 251 | 9.7 | 64 |
|  | 1 mM glutamine | 34 | 1.0 | 0 |
| MalE(T36C/S352C) | no ligand | 503 | 10.9 | 0 |
|  | 1 mM maltotetraose | 32 | 1.2 | 0 |
| | 1.2 μM maltose | 37 | 1.6 | 378 |
|  | 400 nM maltotriose | 144 | 7.2 | 968 |
|  | 450 nM maltotetraose | 70 | 1.8 | 345 |
| | 2 μM maltoheptaose | 61 | 4.9 | 591 |
| | 2 μM maltooctaose | 50 | 2.2 | 491 |
| | 4 μM maltodecaose | 75 | 4.9 | 257 |
| MalE(T36C/S352C/A96W/I329W) | 10 nM maltose | 20 | 20.4 | 30 |
| PsaA(76C/K237C) | no ligand | 254 | 26.3 | 0 |
| | 200 μM Mn<sup>2+</sup> | 35 | 5.8 | 0 |

\* Fluorescence trajectories of 2 independent labelled proteins samples where pooled and analyzed.

**Figure 3.**
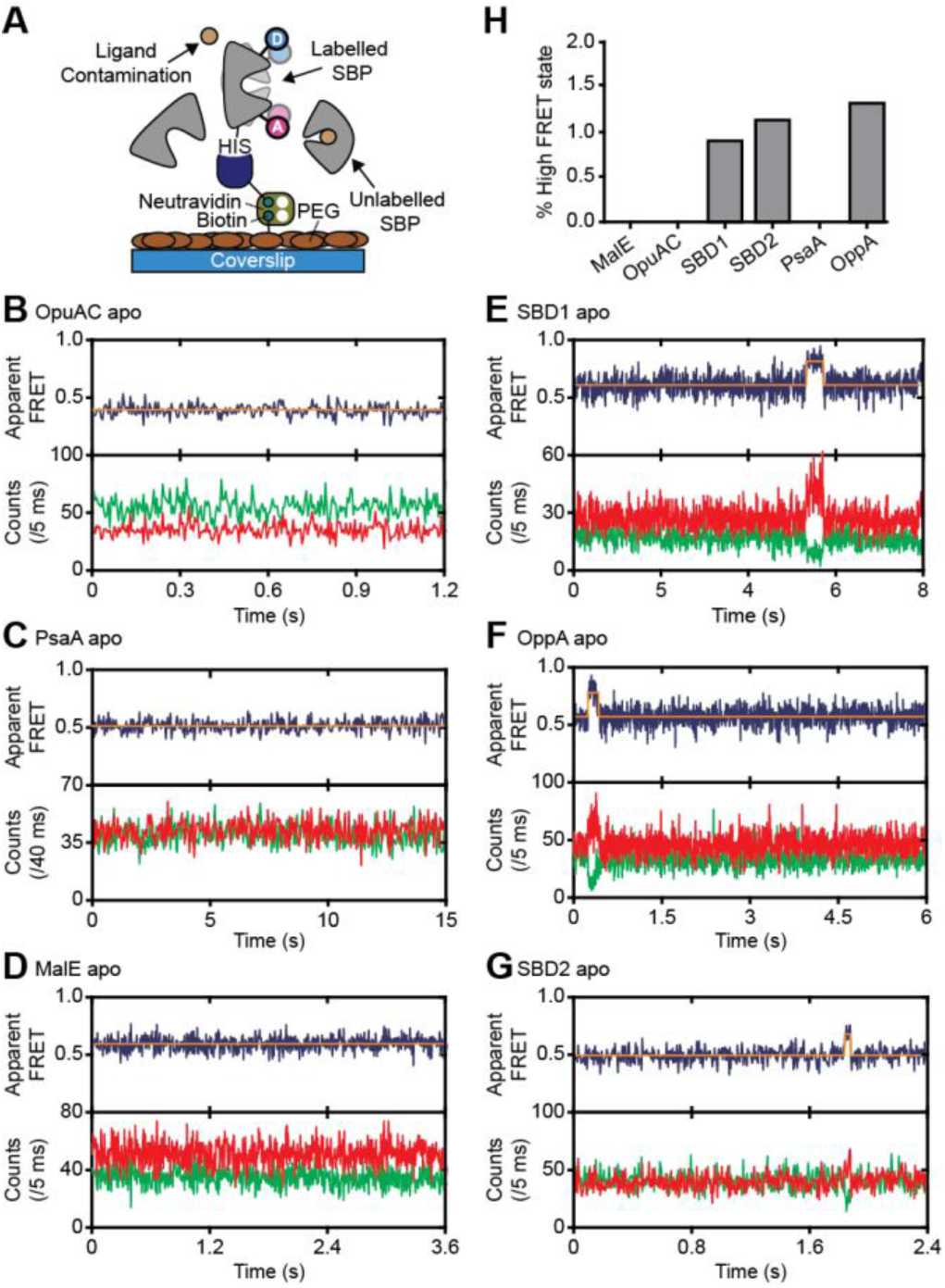
Rare conformational states of ligand-free SBPs. (**A**) Schematic of the experimental strategy to study the conformational dynamics of ligand-free SBPs. Representative fluorescence trajectories of OpuAC(V360C/N423C) (**B**), PsaA(V76C/K237C) (**C**), MalE(T36C/S352C) (**D**), SBD1(T159C/G87C) (**E**), OppA(A209C/S441C) (**F**) and SBD2(T369C/S451) (**G**) in the absence of substrate. 20 μM of unlabelled protein or 1 mM EDTA (for PsaA) was added to scavenge any ligand contaminations. In all fluorescence trajectories presented in the figure: top panel shows calculated apparent FRET efficiency (blue) from the donor (green) and acceptor (red) photon counts as shown in the bottom panels. Orange lines indicate average apparent FRET efficiency value or most probable state-trajectory of the Hidden Markov Model (HMM). Statistics can be found in Table 4. (**H**) Percentage of time a SBP is in the high FRET efficiency state. Statistics can be found in Table 4

### How do non-transported substrates influence the SBP conformation?

Ensemble FRET measurements using all proteinogenic amino acids and citruline were performed to obtain full insight into substrate specificity of SBD1 and SBD2 of GlnPQ. We find that asparagine, glutamine and histidine elicit a FRET change in SBD1, and glutamine in SBD2 (**Figure 4 – figure supplements 1**); glutamate triggers a change in SBD2 at low pH, that is, when a substantial fraction of glutamic acid is present. No other amino acid affected the apparent FRET efficiency. Arginine and lysine, however, competitively inhibit the conformational changes induced by asparagine binding to SBD1 and glutamine binding to SBD2 (**Figure 4 – figure supplements 2**). Uptake experiments in whole cells and in proteoliposomes show that histidine, lysine and arginine are not transported by GlnPQ, but these amino acids can inhibit the uptake of glutamine (via SBD1 and SBD2) and asparagine (via SBD1) (**Figure 4A-C**). Thus, some amino acids interact with the SBPs of GlnPQ but fail to trigger transport. Similar ligands have been identified for MalE, OpuAC and PsaA^16, 38–40^, and we refer to these as non-cognate substrates. We then used smFRET to test whether or not ligand-induced SBP conformational changes allow discriminating cognate from non-cognate substrates.

**Figure 4.**
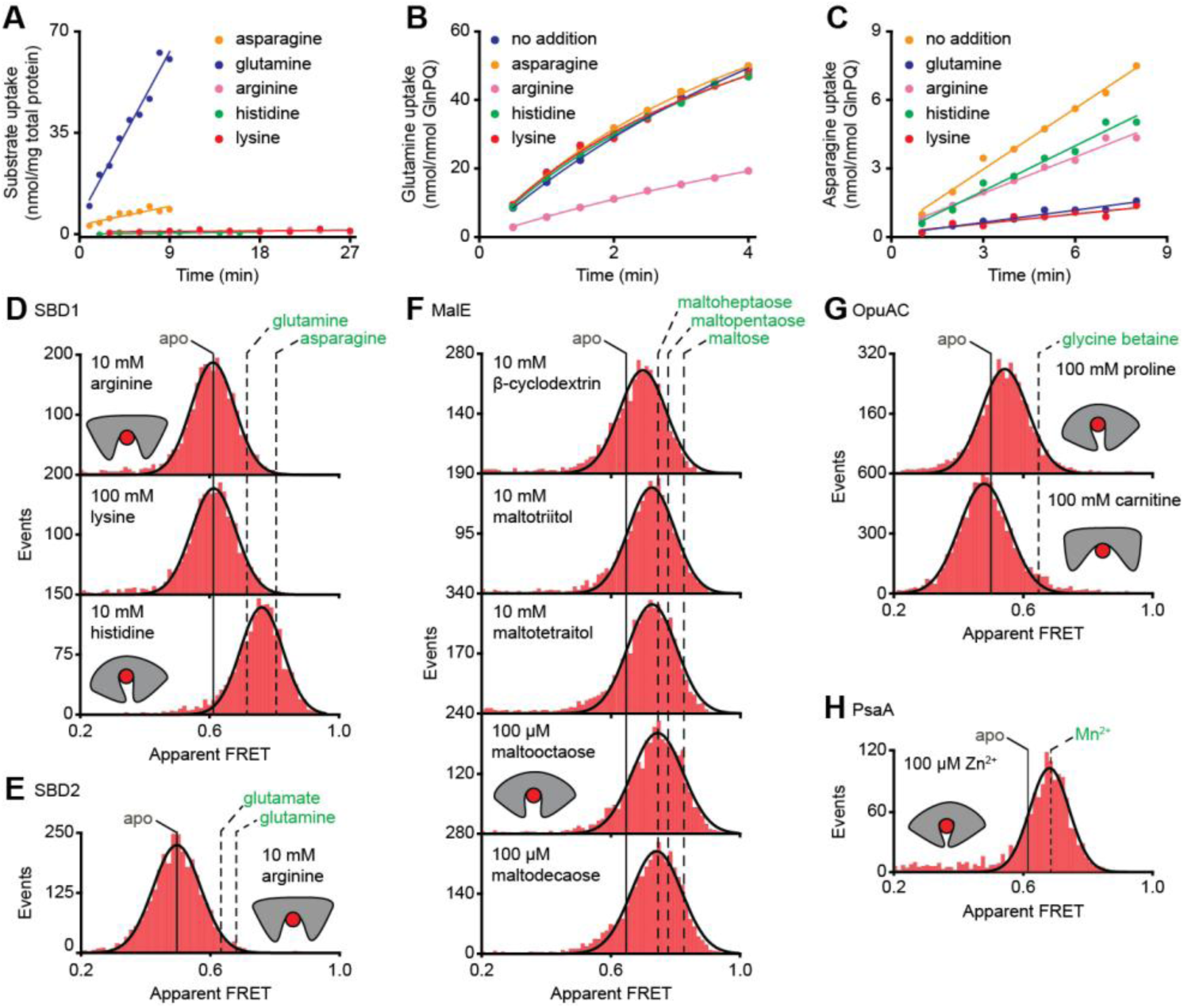
Substrate-specificity of GlnPQ and SBP conformations induced by non-cognate substrates. (**A**) Time-dependent uptake [^14^C]-asparagine (5 μM), [^14^C]-glutamine (5 μM), [^14^C]-arginine (100 μM), [^14^C]-histidine (100 μM) and [^3^H]-lysine (100 μM) by GlnPQ in *L. lactis* GKW9000 complemented *in trans* with a plasmid for expressing GlnPQ; the final amino acid concentrations are indicated between brackets. Points are the data and the solid line a hyperbolic fit. Time-dependent uptake of glutamine (**B**) and asparagine (**C**) in proteoliposomes reconstituted with purified GlnPQ (see Methods section). The final concentrations of [^14^C]-glutamine and [^14^C]-asparagine was 5 μM, respectively; the amino acids indicated in the panel were added at a concentration of 5 mM. Solution-based apparent FRET efficiency histogram of SBD1(T159C/G87C) (**D**), SBD2(T369C/S451) (**E**), MalE(T36C/S352C) (**F**), OpuAC(V360C/N423C) (**G**) and PsaA(V76C/K237C) (**H**) in the presence of non-cognate (red bars) substrates as indicated. Bars are experimental data and solid line a Gaussian fit. The 95% confidence interval for the distribution mean is shown in Table 3. The interval center is indicated by vertical lines (solid and dashed) for the indicated condition.

For most non-cognate substrates, we observe at saturating concentrations that the mean FRET efficiencies are altered compared to the ligand-free conditions (**Figure 4D-H**; **Table 3**). This shows that, similar to cognate ligands (**Figure 3B-G**), non-cognate ligand binding is coupled to SBP conformational changes. However, this is not generally true, as the binding of the non-cognate substrates, i.e., arginine or lysine for SBD1 and arginine for SBD2 do not alter the FRET efficiency histograms (**Figure 4D-E**), suggesting that these ligands bind in the open conformation of the SBP and do not trigger a conformational change.

Further analysis of the non-cognate ligand-induced conformational changes reveals states that vary, from larger opening (carnitine-OpuAC, **Figure 4G**), to partial (histidine-SBD1, **Figure 4D**; various maltodextrin-MalE complexes, **Figure 4F**; proline-OpuAC, **Figure 4G**) or full closing (Zn^2+^-PsaA, **Figure 4H**) of the SBP relative to the ligand-free state of the corresponding protein. The data of full closing by Zn^2+^ (non-cognate) and Mn^2+^ (cognate) were confirmed by examining different inter-dye positions in PsaA (**Table 3**) and are in line with prior crystallographic analyses^40, 50^. Noteworthy, the non-cognate substrate histidine and the cognate substrate glutamine induce both partial closing of SBD1 (**Figure 4D**). However, histidine elicited a larger FRET shift in SBD1 than glutamine, but smaller than the cognate substrate asparagine, which induces full closing (**Figure 4D**). On the other hand, the FRET shift induced with certain non-cognate ligands in MalE (β-cyclodextrin, maltotriitol and maltotetraitol) and OpuAC (proline) are smaller (or similar; *vide infra*) than with their cognate ligands (**Figure 4F**-**G**). Intriguingly, the data also suggest that the partially closed SBP-ligand complexes of MalE formed with the non-cognate substrates maltooctaose or maltodecaose are similar to that of the cognate substrate maltoheptaose (**Figure 4F**). Again, this result was confirmed by examining different inter-dye positions in MalE (**Table 3**).

In summary, similar to cognate substrates, non-cognate substrates do not induce a single unique ligand-bound SBP state, and solely from the degree of SBP closing a translocator cannot readily discriminate cognate from non-cognates substrates. Notable exceptions are the substrates that do not induce closing and keep the SBP in the open state or even to a more extended state. This raises fundamental questions as to the mechanistic basis for how certain non-cognate substrates are still excluded from import.

### Altered SBP opening renders PsaA permissive for non-cognate ligand transport

The inability of certain substrates to be transported, while they appear to induce SBP conformations that are similar to those associated with cognate substrates, was observed for MalE (**Figure 4F**) and PsaA (**Figure 4H**). First, this was investigated further for PsaA. Upon addition of 1 mM EDTA to PsaA-Mn^2+^, lower FRET efficiencies are instantaneously recorded (**Figure 5A**), indicating that the lifetime of the closed PsaA-Mn^2+^ conformation is shorter than a few seconds. By contrast, Zn^2+^ kept PsaA closed, irrespective of the duration of the EDTA treatment (up to 15 min) (**Figure 5B**). Irreversible and reversible binding of these metals was shown previously^51^, which can now be explained by the extremely slow and fast opening of PsaA in the presence of Mn^2+^and Zn^2+^, respectively. The slow opening of PsaA may explain why Zn^2+^ is not transported by PsaBCA, but it is also possible that the TMDs controls the transport specificity^20, 21^. To discriminate between these two scenarios, we examined the impact of altered SBP dynamics on the transport activity of PsaBC. We substituted an aspartate in the binding site with asparagine (D280N), which has previously been shown to perturb the stability of the Zn^2+^-bound SBP^51^. Analysis of PsaA and PsaA(D280N), at saturating Zn^2+^ concentrations, revealed similar FRET efficiency histograms for the two proteins (**Figure 5C**, **Table 3**). However, in contrast to the Zn^2+^-PsaA complex, opening of the PsaA(D280N) complex renders Zn^2+^ accessible to EDTA, similar to the cognate ligand Mn^2+^ (**Figure 5A**,**C**). The ability of PsaA(D280N) to open and release Zn^2+^ was then assessed by measuring the cellular accumulation of Zn^2+^ within *Streptococcus pneumoniae*, the host organism. This was achieved by replacement of the *psaA* gene with the D280N mutant allele (Ω*psaA*D280N) in a strain permissive for Zn^2+^ accumulation, i.e. incapable of Zn^2+^ efflux due to deletion of the exporter CzcD (Ω*psaA*_D280N_Δ*czcD*)^52^. Our data show that cellular Zn^2+^ accumulation increases in the strain expressing PsaBC with PsaA(D280N) but not with wild-type PsaA (**Figure 5D**). These results demonstrate that the altered conformational dynamics of the PsaA derivative renders ligand release permissive for transport of non-cognate Zn^2+^ ions. The data also show that translocator activity is not directly influenced by the nature of the metal ion released by PsaA. Collectively, our findings show that transport specificity of PsaBCA is dictated by the opening kinetics of PsaA.

**Figure 5.**
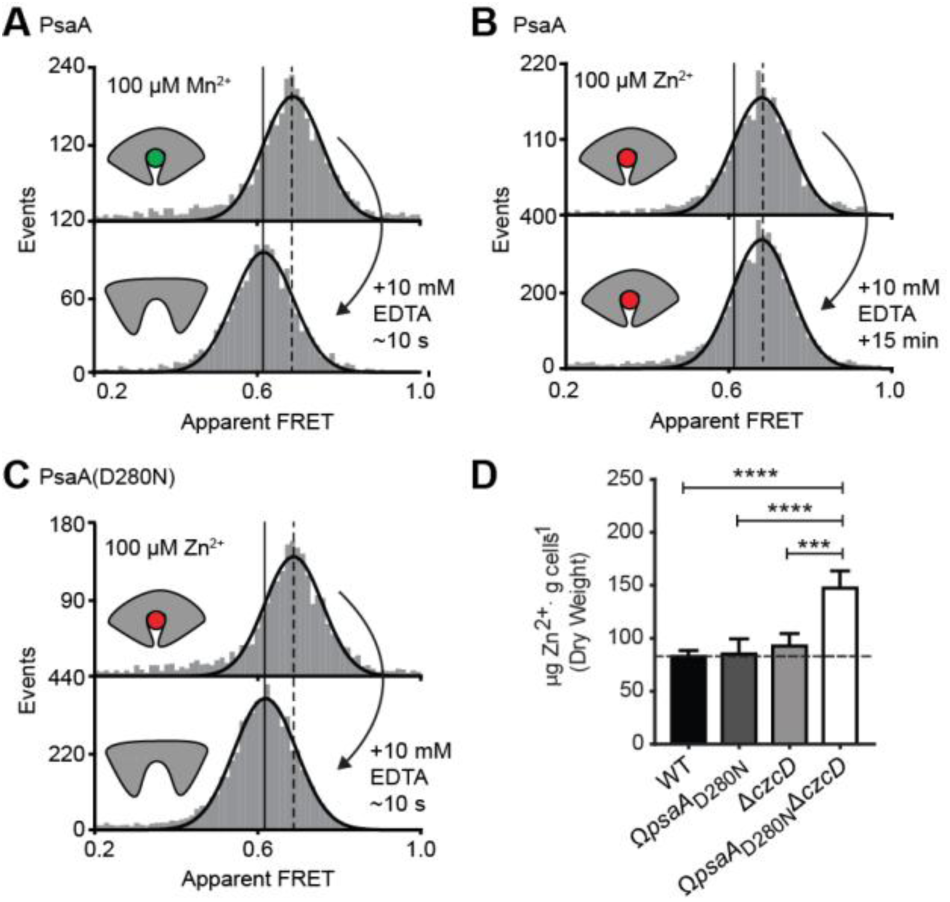
Opening transition in PsaA dictates transport specificity. Solution-based apparent FRET efficiency histograms of PsaA(V76C/K237C) in the presence of Mn^2+^ (**A**) or Zn^2+^ (**B**) and PsaA(D280N) in the presence of Zn^2+^ (**C**) upon addition of 10 mM EDTA and incubated for the indicated duration. Bars are experimental data and the solid line a Gaussian fit. The 95% confidence interval for the mean of the Gaussian distribution can be found in Table 3, and the interval centre is indicated by vertical lines (solid, metal-free and dashed, metal-bound). (**D**) Whole cell Zn^2+^ accumulation of *S. pneumoniae* D39 and mutant strains in CDM supplemented with 50 µM ZnSO_4_as determined by ICP-MS. Data correspond to mean ± s.d. μg Zn^2+^.g^−1^dry cell weight from three independent biological experiments. Statistical significance was determined by one-way ANOVA with Tukey post-test (\*\*\**P*<0.005 and \*\*\*\**P*<0.0001).

### MalE conformational dynamics with cognate and non-cognate substrates

Next, we determined the conformational dynamics of MalE induced by maltoheptaose, maltooctaose and maltodecaose. Similar to Zn^2+^ and Mn^2+^ in PsaA (**Figure 4H**), these substrates appear to induce similar MalE conformations (**Figure 4F**), but only maltoheptaose is transported^39^. Measurements on individual surface-tethered MalE proteins, in the presence of maltoheptaose, maltooctaose or maltodecaose, showed frequent switching between low and higher FRET states, corresponding to opening and closing of the protein (**Figure 6A-D**). The mean lifetime of the ligand-bound conformations, e.g. the mean lifetime of the higher FRET states, are 328 ± 8 ms for cognate maltoheptaose and 319 ± 12 ms and 341 ± 8 ms for non-cognate maltooctaose and maltodecaose, respectively (mean ± s.e.m.; **Figure 6A**, **Figure 6 – figure supplements 6**). So, contrary to PsaA-Zn^2+^ (**Figure 5**), a slow opening of MalE and inefficient ligand release kinetics cannot explain why maltooctaose and maltodecaose are not transported; the average lifetimes with maltooctaose or maltodecaose are not significantly different from that with maltoheptaose (*P*=0.68, one-way analysis of variance (ANOVA); **Figure 6A**). Most likely, the failure of the maltose system to transport maltooctaose and maltodecaose originates in the dimensions of the substrate cavity within the translocator domain of MalFGK_2_^20^.

**Figure 6.**
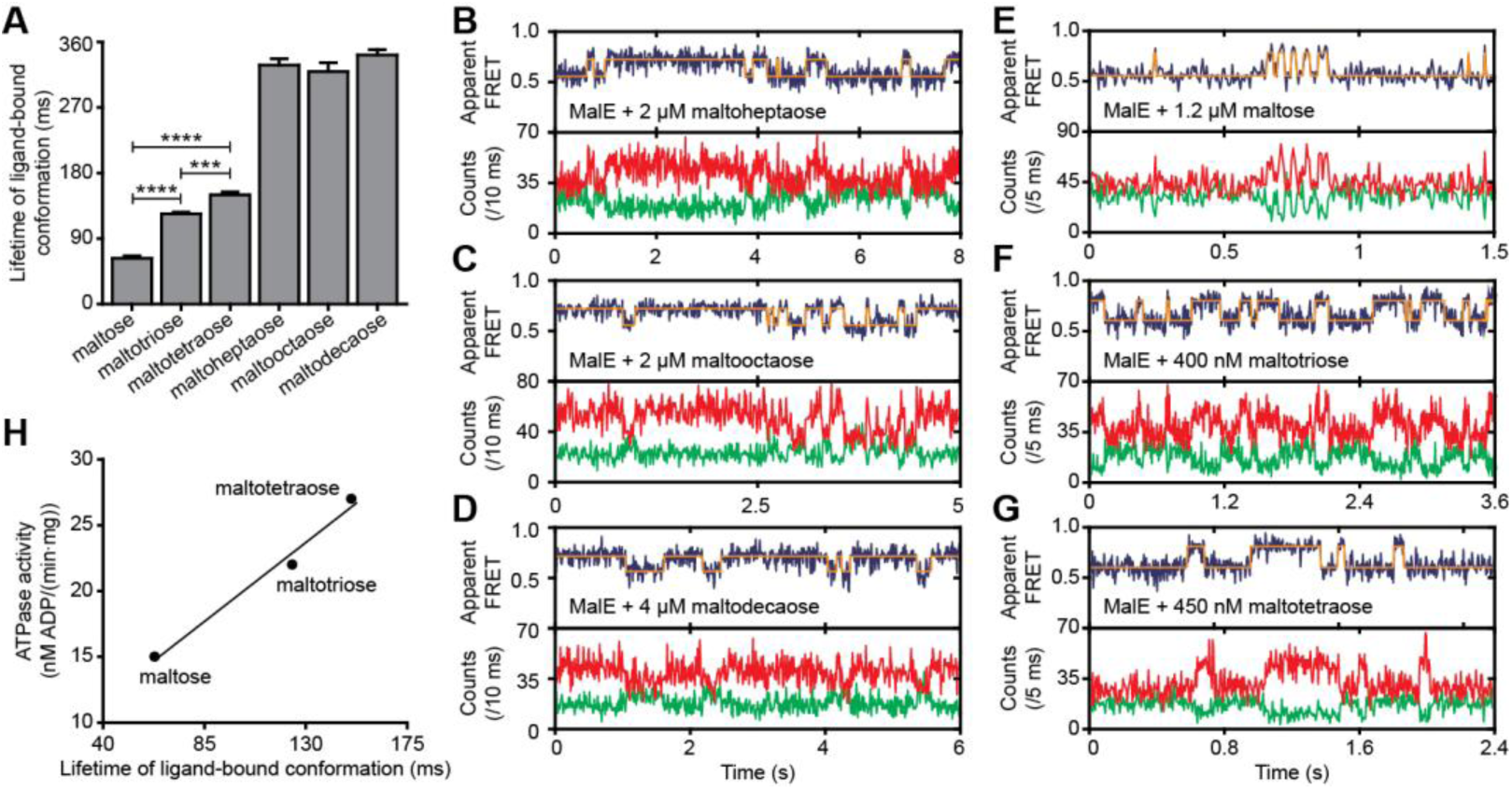
Lifetime of MalE ligand-bound conformations and relation to activity. (**A**) Mean lifetime of the ligand-bound conformations of MalE, obtained from all single-molecule fluorescence trajectories in the presence of different maltodextrins as indicated. Data corresponds to mean ± s.e.m.. Histogram of the data are shown in Figure 6 – figure supplement 1. The statistical significance of the differences in the mean data was determined by two-tailed unpaired *t*-tests (\*\*\**P*<0.005 and \*\*\*\**P*<0.0001). (**B**, **C**, **D**, **E**, **F** and **G**) Representative fluorescence trajectories of MalE(T36C/S352C) in the presence of different substrates as indicated. In all fluorescence trajectories presented: top panel shows calculated apparent FRET efficiency (blue) from the donor (green) and acceptor (red) photon counts as shown in the bottom panels. Most probable state-trajectory of the Hidden Markov Model (HMM) is shown (orange). (**H**) Published ATPase activity^16^ linked to the lifetime of the closed MalE conformation induced by transport of different cognate substrates as indicated. Points are the data and the solid line a simple linear regression fit.

### Translocator/SBP interplay determines the rate of transport

Finally, we sought to elucidate the mechanistic basis for how substrate preference arises in the maltose system and to what degree the translocator contributes to this process. First, we investigated how the MalE conformational dynamics influences the transport rate of the substrate maltose. For this we used the hinge-mutant variant MalE(A96W/I329W) that has different conformational dynamics compared to the wild-type protein (**Figure 6E**; **Figure 6 – figure supplements 6A-B**)^32^. The mutations are believed to not affect SBP-translocator interactions since they are situated on the opposite side of the interaction surface of the SBP^44, 53^.

In the presence of saturating concentrations of maltose the FRET efficiency distributions of MalE and MalE(A96W/I329W) are indistinguishable. This could be confirmed by two different inter-dye positions in each protein (**Figure 6 – figure supplements 6C**). Therefore, changes in the rate of maltose transport unlikely arise from differences in SBP docking onto the TMD, since similar SBP conformations are involved. Nonetheless, cellular growth and the maltose-induced ATPase activity are reduced for MalE(A96W/I329W)^53, 54^. Analysis of the mean lifetime of the closed conformation of MalE(A96W/I329W) shows that ligand release is three orders of magnitude slower than in the wild-type protein [63 ± 6 ms (mean ± s.e.m.) in MalE *versus* 94 ± 16 s (mean ± s.e.m.) in MalE(A96W/I329W); **Figure 6A**; **Figure 6 – figure supplements 6B**]. These observations suggest that the maltose-stimulated cellular growth and ATPase activity are reduced due to the slower opening of MalE(A96W/I329W) compared to wildtype MalE. All this is in line with the observation that PsaA(D280N) opens fast, allowing transport of the Zn^2+^ to occur, whereas in wildtype PsaA the opening of PsaA after Zn^2+^-binding is (extremely) slow and transport does not occur (**Figure 5B-D**).

We then investigated the relationship between maltodextrin-specific lifetimes of the MalE closed conformations and published transport rates or ATPase activities of the full transport system^16^. Here, we focused on the cognate substrates maltose, maltotriose and maltotetraose since crystallographic^15^ and smFRET analysis (**Figure 2 – figure supplements 2A**; **Table 3**) suggest that these ligands induce similar MalE conformations. The average lifetime of the closed conformation with maltose, maltotriose and maltotetraose are 63 ± 6, 124 ± 4, and 150 ± 8 ms (mean ± s.e.m.), respectively (**Figure 6A**; **Figure 6E-G**; **Figure 6 – figure supplements 1**). Thus, these lifetimes correlate positively with their stimulation of the ATPase activity (**Figure 6H**)^16^. A positive relation also exists between the lifetimes with maltose and maltotetraose (63 ± 6 and 150 ± 8 ms; mean ± s.e.m.) and their corresponding transport rates (transport of maltotetraose is ∼1.5-fold higher than of maltose)^16^. The observation that some maltodextrins induce a faster opening of MalE (short lifetime), while their corresponding transport and/or stimulation of ATP hydrolysis are slower, implies an involvement of the translocator MalFGK_2_ in causing the variability in the transport rate of these maltodextrins.

## DISCUSSION

Prokaryotes occupy diverse ecological niches within terrestrial ecosystems. Irrespective of the niche, their viability depends on selective acquisition of nutrients from the extracellular environment. However, the diversity of the external milieu poses a fundamental challenge for how acquisition of specific compounds can be achieved within the constraints of the chemical selectivity conferred by their import pathways. Numerous studies on SBPs associated with ABC importers have established that these proteins share a common architecture with a well-defined high-affinity ligand-binding site and have the ability to adopt distinct ligand-free and -bound conformations, i.e. open and closed, respectively^2, 7, 8^. Building on this knowledge, we investigated the relationship between SBP conformational dynamics, SBP-ligand interactions and substrate transport.

The general view of SBP conformational changes serving as a binary switch to communicate transport competency may hold for some SBPs, such as OppA (**Figure 2 – figure supplements 2B**), while others employ multiple distinct ligand-bound conformations (**Figure 2D-F**; **Figure 4D-G**). To our knowledge, such extreme conformational plasticity of SBPs has not been observed before. MalE shows a remarkable structural flexibility of at least six different ligand-bound conformations (**Figure 2D**; **Figure 4F**). SBD1 (**Figure 2E**; **Figure 4D**) can sample at least four distinct ligand-bound conformations and SBD2 (**Figure 2F**; **Figure 4E**) and OpuAC (**Figure 2B**; **Figure 4G**) at least three. Moreover, MalE, SBD1 and SBD2 have multiple distinct ligand-bound conformations that can all interact with the translocator, as they all facilitate substrate import (’multiple conformations activate transport’ in **Figure 7**; **Figure 2D-F**). Thus, a productive SBP-translocator interaction in Type I ABC importers can be accomplished without relying on strict structural requirements for docking. This generalization may not apply to all Type I ABC importers since in the Opp importer the translocator might only interact with a unique closed conformation of the SBP OppA (**Figure 2 – figure supplements 2B**), and Opp has no measurable affinity for its open ligand-free conformation^46^.

**Figure 7.**
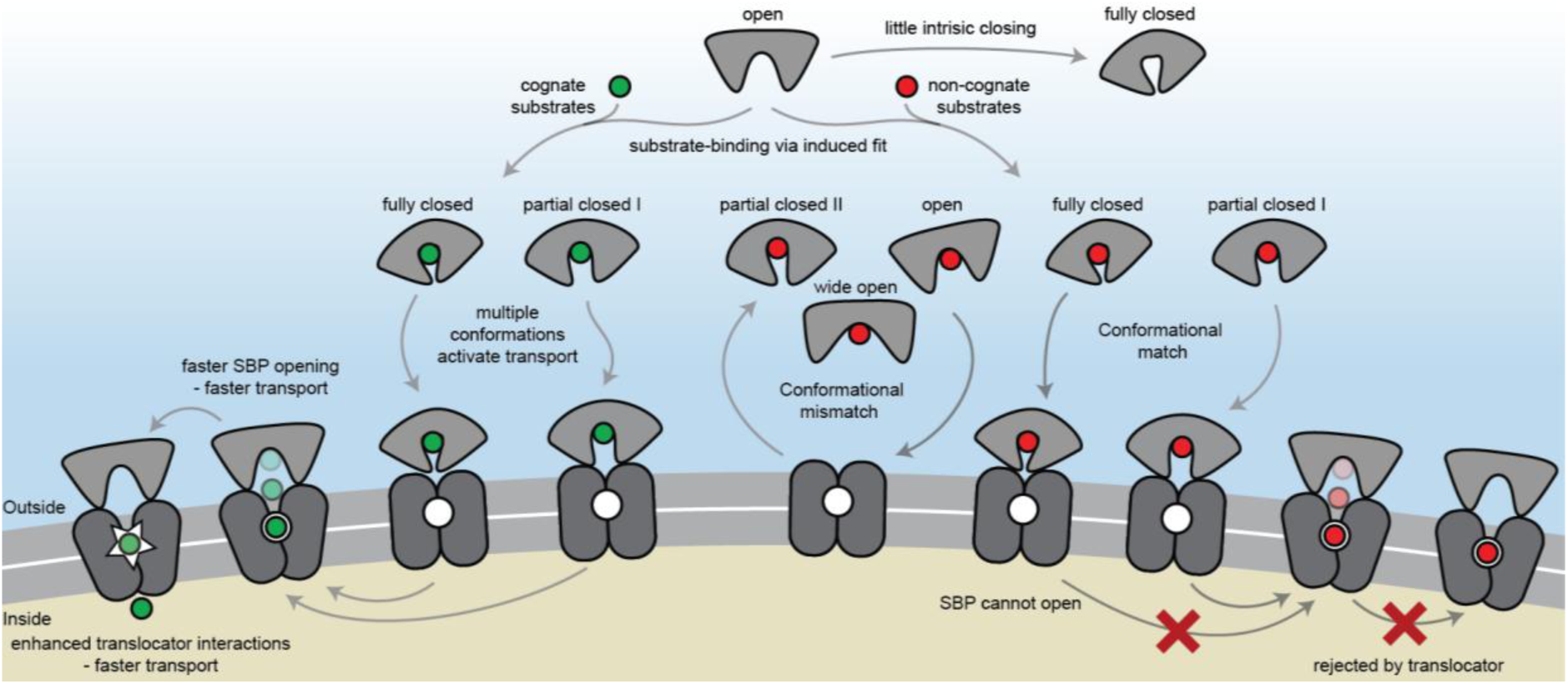
The conformational changes and dynamics of SBPs and the regulation of transport. Schematic summarizing the plasticity of ligand binding and solute import via ABC importers. Intrinsic closing of an SBP is a rare event or absent at all in some SBPs (‘little intrinsic closing’). Ligands are bound via induced fit (‘ligand-binding via induced fit’). SBPs can acquire one or more conformations that can activate transport (‘multiple conformations activate transport’). Variations in cognate substrate transport are caused by: (i) openings rate of the SBP and substrate transfer to the translocator (‘faster SBP opening – faster transport’) and (ii) substrate-dependent translocator interactions (‘enhanced translocator interactions – faster transport’). Although SBPs can acquire a conformation that activates transport (‘conformational match’), transport still fails when: (i) the SBP has no affinity for the translocator and/or cannot make the allosteric interaction with the translocator (‘conformational mismatch’); (ii) the SBP cannot open and release the substrate to the translocator (‘SBP cannot open’); or (iii) due to the specificity of the translocator (‘rejected by translocator’).

Exclusion of non-cognate substrates is also a critical biological function for SBPs. Our work has uncovered a hitherto unappreciated complexity in protein-ligand interactions and how this is coupled to regulation of substrate import. Similar to transport, exclusion of non-cognate ligands might be achieved by multiple distinct mechanisms. We have shown that although multiple SBP conformations can activate transport (**Figure 2D-F**), not all SBP conformational states appear to provide the signal to facilitate transport. For example, the binding of certain non-cognate ligands induces a conformational change in SBD1 (**Figure 4D**), MalE (**Figure 4F**) and OpuAC (**Figure 4G**) that are distinct from those that facilitate transport. However, non-cognate substrate binding is not always coupled to an SBP conformational change, as observed for the binding of arginine or lysine to SBD1 and arginine to SBD2 (**Figure 4D-E**). These observations provide a general explanation on how substrate import can fail in Type I ABC importers, which would be due to the SBP-ligand complex assuming a conformation that cannot initiate allosteric interactions with the translocator (‘conformational mismatch’ in **Figure 7**). A similar hypothesis was put forward based on the observation that binding of β-cyclodextrin fails to fully close MalE^17–19^. However, the sole observation of partial closing of MalE cannot explain why transport of β-cyclodextrin fails, as we here show that also cognate maltodextrins are able to induce partial closing of MalE (**Figure 2D**).

By contrast, in the Mn^2+^ transporter PsaBCA, a different mechanism is used. In PsaA, the binding site composition of the SBP precludes the ability of the protein to exclude the non-cognate substrate Zn^2+^ from interacting. As a consequence, both metals bind and trigger formation of similar PsaA conformations (‘conformational match’ in **Figure 7**; **Figure 4H**)^40, 50^. Despite this, the two ions have starkly different conformational dynamics, with Zn^2+^ forming a highly stable closed conformation, such that it cannot open and release the substrate to its translocator (‘SBP cannot open’ in **Figure 7**; **Figure 5**). By altering the binding site interactions between PsaA and Zn^2+^, opening is faster and transport of the metal ion can occur (**Figure 5B-D**). Similar observations were made for GlnPQ^29, 55^ and MalE (**Figure 6E**, **Figure 6 – figure supplement 2A**), in which a slower/faster opening of the SBP resulted in a decrease/increase in the corresponding transport of the substrate or ATP hydrolysis rate (‘faster SBP opening – faster transport’ in **Figure 7**). We therefore conclude that for ligands that induce highly stabilized SBP-substrate conformations, which require more energy (thermal or ATP-dependent) to open, transport becomes slower or is abrogated. Based on these findings, we infer that biological selectivity in ABC importers is largely achieved via a combination of ligand release kinetics and its influence on the conformational state of the SBP. This provides a mechanism to facilitate the import of selective substrates, while excluding other compounds. However, our data also implicate a role for the translocator in contributing to the substrate specificity of ABC importers, consistent with previous studies^20, 21, 48, 56^.

The presence of a substrate binding site in the translocator of the maltose system is well established^20, 48^, although its role, if any, in influencing the rate of transport of maltodextrins is yet unknown. The average time required for the different maltodextrin-MalE complexes to open, correlates positively with the transport and ATP hydrolysis rate (**Figure 6H**)^16^. This implies that the substrate, after it has been transferred from MalE to the translocator, acts as a trigger for subsequent steps, for example, the stimulation of ATP hydrolysis and/or Pi and ADP release (‘enhanced translocator interactions – faster transport’ in **Figure 7**). The positive correlation implies that some maltodextrins trigger these steps more efficient than others, thereby overcoming the slower opening of MalE, and leading to a preferred uptake of certain maltodextrins over others. Further, analysis of the non-cognate substrates maltooctaose and maltodecaose showed that these were bound reversibly by MalE (**Figure 6A**) and can induce a conformation similar that to that of the cognate ligand maltoheptaose (‘conformational match’ in **Figure 7**; **Figure 4F**). We speculate that the failure of the maltose system to transport maltooctaose and maltodecaose most likely arises from the dimensions of the substrate cavity within the MalFGK2^20^ translocator (‘rejected by translocator’ in **Figure 7**), rather than failure of MalE to close and release the substrate.

The presence of two consecutive binding pockets, one in the SBP and one in the translocator, in at least some ABC importers could indicate that specificity of transport occurs through a proofreading mechanism in a manner analogous to aminoacyl-tRNA synthetases and DNA polymerase^57, 58^. In such a mechanism, a substrate can be rejected even if it has been bound by the SBP. Although we show that intrinsic closing is a rare event (‘little intrinsic closing’ in **Figure 7**; data in **Figure 3**), it might influence transport in a cellular context where the ratio between SBP and translocator can be high^59^. Moreover, other fast (µs-ms) and short-range conformational changes might be present as shown by NMR analysis on MalE^27^. We speculate that in Type I ABC importers the wasteful conversion of chemical energy is prevented by a proofreading mechanism, as any thermally driven closing event would not be able to initiate the translocation cycle, as the substrate is absent. In accordance, ATP hydrolysis and transport are tightly coupled in the Type I importer GlnPQ^60^ that, based on the crystal structure of the homologous Art(QM)_2_ ^21^, contains an internal binding pocket located within the TMDs. By contrast, high futile hydrolysis of ATP in the Type II BtuCDF^61^ appears to correlate with the lack of a defined binding pocket inside the TMDs.

## METHODS

### Gene expression and SBP purification

N-terminal extension of the soluble SBPs with a Hisx tag (His_10_PsaA, His_10_SBD1, His_10_SBD2, His_10_OppA and His_6_OpuAC) were expressed and purified as previously described^29, 38, 43, 51^. The *mal*E gene (UniProtKB-P0AEX9) was isolated from the genome of *Escherichia coli* K12. The primers were designed to exclude the signal peptide (amino acids 1-26). Primers introduced *Nde*I and *Hind*III restriction sites, and the gene product was sub-cloned in the pET20b vector (Novagen, EMD Millipore). Protein derivatives having the cysteine or other point mutations were constructed using QuikChange mutagenesis^62^ and Megaprimer PCR mutagenesis^63^ protocols. Primers are indicated in Table 5. His_6_MalE was over-expressed in *E. coli* BL21 DE3 cells (*F–ompT gal dcm lon hsdSB*(*r*_*B*_*–m*_*B*_) λ(DE3 [*lacI lacUV5-T7p07 ind1 sam7 nin5*]) [malB+]K-12(λS)). Cells harbouring plasmids expressing the MalE wild-type and derivatives were grown at 30°C until an optical density (OD_600_) of 0.5 was reached. Protein expression was then induced by addition of 0.25 mM isopropyl β-D-1-thiogalactopyranoside (IPTG). After 2 hours induction cells were harvested. DNase 500 ug/ml (Merck) was added and passed twice through a French pressure cell at 1,500 psi and 2mM phenylmethylsulfonyl fluoride (PMSF) was added to inhibit proteases. The soluble supernatant was isolated by centrifugation at 50,000 × *g* for 30 min at 4 °C. The soluble material was then purified and loaded on Ni^2+−^sepharose resin (GE Healthcare) in 50 mM Tris-HCl, pH 8.0; 1 M KCl, 10 % glycerol; 10 mM imidazole, 1 mM dithiothreitol (DTT). The immobilized proteins were washed (50 mM Tris-HCl, pH 8.0; 50 mM KCl, 10 % glycerol; 10 mM Imidazole; 1 mM DTT; plus 50 mM Tris-HCl, pH=8; 1 M KCl, 10 % glycerol; 30 mM imidazole; 1 mM DTT sequentially) and then eluted (50 mM Tris-HCl, pH 8.0, 50 mM KCl, 10 % glycerol; 300 mM imidazole; 1 mM DTT). Protein fractions were pooled (supplemented with 5 mM EDTA, 10 mM DTT), concentrated (10.000 MWCO Amicon; Merck-Millipore), dialyzed against 100-1000 volumes of buffer (50 mM Tris-HCl, pH 8.0; 50 mM KCl, 50% glycerol; 10 mM DTT), aliquoted and stored at −20°C until required.

**Table 5.**
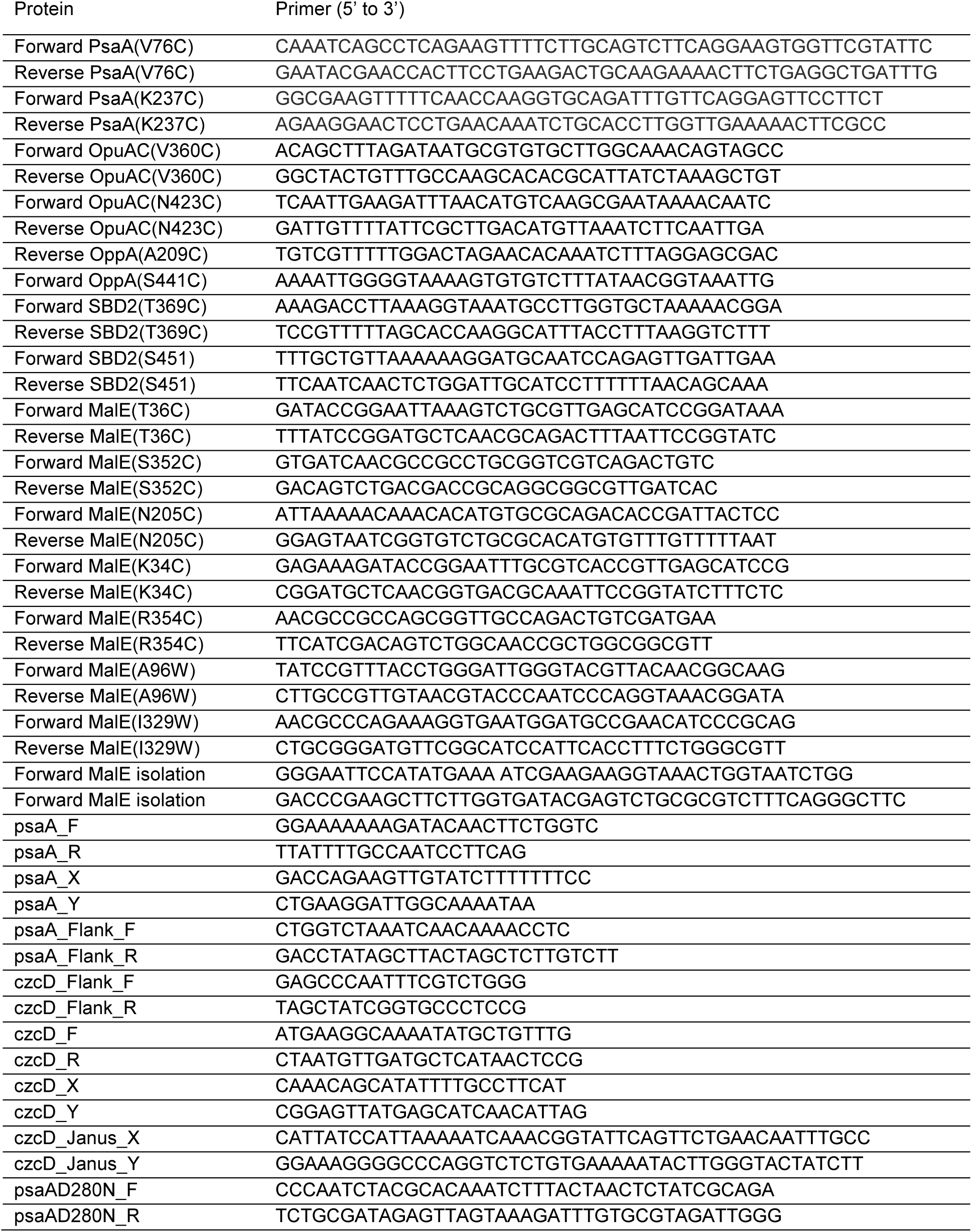
Primers used in this study.

### Uptake experiments in whole cells

*Lactococcus lactis* GKW9000 carrying pNZglnPQhis or derivatives was cultivated semi-anaerobically at 30 ^o^C in M17 (Oxoid) medium supplemented with 1 % (w/v) glucose and 5 μg/ml chloramphenicol. For uptake experiments cells were grown in GM17 to an OD_600_ of 0.4, induced for 1 hour with 0.01 % of culture supernatant of the nisin A-producing strain NZ9700 and harvested by centrifugation for 10 min at 4000 × *g*; the final nisin A concentration is ∼1 ng/ml. After washing twice with 10 mM PIPES-KOH, 80 mM KCl, pH 6.0, the cells were resuspended to OD_600_ = 50 in the same buffer. Uptake experiments were performed at 0.1 – 0.5 mg/ml total protein in 30 mM PIPES-KOH, 30 mM MES-KOH, 30 mM HEPES-KOH (pH 6.0). Before starting the transport assays, the cells were equilibrated and energized at 30°C for 3 min in the presence of 10 mM glucose plus 5 mM MgCl_2_. After 3 min, the uptake reaction was started by addition of either [^14^C]-glutamine, [^14^C]-histidine, [^14^C]-arginine, [^14^C]-lysine (all from Perkin Elmer) or [^3^H]-asparagine (ARC); the specific radioactivity was adjusted for each experiment (amino acid concentration) to obtain sufficient signal above background; the final amino acid concentrations are indicated in the figure legends. At given time intervals, samples were taken and diluted into 2 ml ice-cold 100 mM LiCl. The samples were rapidly filtered through 0.45 µm pore-size cellulose nitrate filters (Amersham) and the filter was washed once with ice-cold 100 mM LiCl. The radioactivity on the filters was determined by liquid scintillation counting.

### Purification and membrane reconstitution of GlnPQ for in vitro transport assays

Membrane vesicles of *Lactococcus lactis* GKW9000 carrying pNZglnPQhis were prepared as described before^60^. For reconstitution into proteoliposomes, 150 mg of total protein in membrane vesicles was solubilized in 50 mM potassium phosphate pH 8.0, 200 mM NaCl, 20% glycerol and 0.5% (w/v) DDM for 30 minutes at 4 °C. The sample was centrifuged (12 min, 300,000*xg*) and the supernatant was collected. Subsequently, GlnPQ was allowed to bind to Ni-Sepharose (1.5 ml bed volume) for 1 hour at 4 °C after addition of 10 mM imidazole. The resin was rinsed with 20 column volumes of wash buffer [50 mM potassium phosphate, pH 8.0, 200 mM NaCl, 20% (v/v) glycerol, 50 mM imidazole plus 0.02% (w/v) DDM]. The protein was eluted with 5 column volumes of elution buffer [50 mM potassium phosphate, pH 8.0, 200 mM NaCl, 10% (w/v) glycerol, 500 mM imidazole plus 0.02% (w/v) DDM]. The purified GlnPQ was used for reconstitution into liposomes composed of egg yolk L-α-phosphatidylcholine and purified *E. coli* lipids (Avanti polar lipids) in a 1:3 ratio (w/w) as described before^64^ with a final protein/lipid ratio of 1:100 (w/w). An ATP regenerating system, consisting of 50 mM potassium phosphate, pH 7.0, creatine kinase (2.4 mg/ml), Na_2_-ATP (10 mM), MgSO_4_ (10 mM), and Na_2_-creatine-phosphate (24 mM) was enclosed in the proteoliposomes by two freeze/thaw cycles, after which the vesicles were stored at −80 °C. On the day of the uptake experiment, the proteoliposomes were extruded 13 times through a polycarbonate filter (200-nm pore size), diluted to 3 ml with 100 mM potassium phosphate, pH 7.0, centrifuged (265,000g for 20 min), and then washed and resuspended in 100 mM potassium phosphate, pH 7.0, to a concentration of 50 mg of lipid/ml.

Uptake in proteoliposomes was measured in 100 mM potassium phosphate, pH 7.0, supplemented with 5 µM of [^14^C]-glutamine or [^3^H]-asparagine. This medium, supplemented with or without unlabelled amino acids (asparagine, arginine, glutamine, histidine or lysine), was incubated at 30 °C for 2 min prior to adding proteoliposomes (kept on ice) to a final concentration of 1-5 mg of lipid/ml. At given time intervals, 40 µl samples were taken and diluted with 2 ml of ice-cold isotonic buffer (100 mM potassium phosphate, pH 7.0). The samples were collected on 0.45-m pore size cellulose nitrate filters and washed twice as described above. After addition of 2 ml Ultima Gold scintillation liquid (PerkinElmer), radioactivity was measured on a Tri-Carb 2800TR (PerkinElmer). A single time-dependent uptake experiment is shown in Figure 4A-C and consistent results were obtained upon repetition with an independent sample preparation.

### Zinc accumulation in whole cells

The *S. pneumoniae* D39 mutant strains Ω*psaA*D280N and Δ*czcD* were constructed using the Janus cassette system^65^. Briefly, the upstream and downstream flanking regions of *psaA* and *czcD* were amplified using primers (Table 5) with complementarity to either *psaA*_D280N_ (Ω*psaA*D280N), generated via site-directed mutagenesis of *psaA* following manufacturer instructions (Agilent), or the Janus cassette (Δ*czcD*) and were joined by overlap extension PCR. These linear fragments were used to replace by homologous recombination *psaA* and *czcD*, respectively, in the chromosome of wild-type and Δ*czcD* strains. For metal accumulation analyses, *S. pneumoniae* strains were grown in a cation-defined semi-synthetic medium (CDM) with casein hydrolysate and 0.5% yeast extract, as described previously^66^. Whole cell metal ion accumulation was determined by inductively coupled plasma-mass spectrometry (ICP-MS) essentially as previously described^52^. Briefly, *S. pneumoniae* strains were inoculated into CDM supplemented with 50 μM ZnSO_4_ at a starting OD_600_ of 0.05 and grown to mid-log phase (OD_600_ = 0.3-0.4) at 37 °C in the presence of 5% CO_2_. Cells were washed by centrifugation 6 times in PBS with 5 mM EDTA, harvested, and desiccated at 95 °C for 18 hrs. Metal ion content was released by treatment with 500 μL of 35% HNO_3_ at 95 °C for 60 min. Metal content was analysed on an Agilent 8900 QQQ ICP-MS^51^.

### Isothermal titration calorimetry (ITC)

Purified OppA was dialyzed overnight against 50 mM Tris-HCl, pH=7.4; 50 mM KCl. ITC experiments were carried by microcalorimetry on a ITC200 calorimeter (MicroCal). The peptide (RPPGFSFR) stock solution (200 μM) was prepared in the dialysis buffer and was stepwise injected (2 μl) into the reaction cell containing 20 μM OppA. All experiments were carried out at 25°C with a mixing rate of 400 rpm. Data were analysed with a one site-binding model using, provided by the MicroCal software (MicroCal).

### Protein labelling for FRET measurements

Surface-exposed and non-conserved positions were chosen for Cys engineering and subsequent labelling, based on X-ray crystal structures of OpuAC (3L6G, 3L6H), SBD1 (4AL9), SBD2 (4KR5, 4KQP), PsaA (3ZK7, 1PSZ), OppA (3FTO, 3RYA) and MalE (1OMP, 1ANF). The proteins used in smFRET experiments were OpuAC(V360C/V423C), SBD1(T159C/G87C), SBD2(T369C/S451C), PsaA(V76C/K237C), PsaA(V76C/K237C/D280N), PsaA(E74C/K237C), OppA(A209C/S441C), MalE(T36C/S352C), MalE(T36C/N205C), MalE(K34C/R354C), MalE(T36C/S352C/A96W/I329W) and MalE(K34C/R354C/A96W/I329W). Unlabelled protein derivatives (20-40 mg/ml) were stored at −20 °C in the appropriate buffer (50 mM Tris-HCl, pH 7.4; 50 mM KCl; 50% glycerol for MalE and OppA; 25 mM Tris-HCl, pH 8.0; 150 mM NaCl; 1 μM EDTA; 50% glycerol for PsaA; 50 mM KPi, pH 7.4; 50 mM KCl; 50% glycerol for OpuAC, SBD1 and SBD2) supplemented by 1 mM Dithiothreitol (DTT, Sigma-Aldrich).

Stochastic labelling was performed with the maleimide derivative of dyes Cy3B (GE Healthcare) and ATTO647N (ATTO-TEC) for OpuAC and MalE; SBD1, SBD2, OppA and PsaA were labelled with Alexa555 and Alexa647 (ThermoFisher). The purified proteins were first treated with 10 mM DTT for 30 min to fully reduce oxidized cysteines. After dilution of the protein sample to a DTT concentration of 1 mM the reduced protein were immobilized on a Ni^2+^-Sepharose resin (GE Healthcare) and washed with ten column volumes of buffer A (50 mM Tris-HCl, pH 7.4; 50 mM KCl for MalE and OppA; 25 mM Tris-HCl, pH 8.0; 150 mM NaCl; 1 μM EDTA for PsaA; 50 mM KPi, pH 7.4; 50 mM KCl for OpuAC, SBD1 and SBD2) to remove the DTT. To make sure that no endogenous ligand was left, for some experiments, and prior to removing the DTT, we unfolded the immobilized-SBPs by treatment with 6 M of urea supplemented with 1 mM DTT and refolded them again by washing with buffer A. The resin was incubated 1-8 hrs at 4 °C with the dyes dissolved in buffer A. To ensure a high labelling efficiency, the dye concentration was ∼20-times higher than the protein concertation. Subsequently, unbound dyes were removed by washing the column with at least twenty column volumes of buffer A. Elution of the proteins was done by supplementing buffer A with 400 mM Imidazole (Sigma-Aldrich). The labelled proteins were further purified by size-exclusion chromatography (Superdex 200, GE Healthcare) using buffer A. Sample composition was assessed by recording the absorbance at 280 nm (protein), 559 nm (donor), and 645 nm (acceptor) to estimate labelling efficiency. For all proteins the labelling efficiency was >90%.

### Fluorescence Anisotropy

To verify that the measurements of apparent FRET efficiency report on inter-probe distances between the donor and acceptor fluorophores, at least one of the fluorophores must be able to rotate freely. To investigate this, we determined the anisotropy values of labelled proteins. The fluorescence intensity was measured on a scanning spectrofluorometer (Jasco FP-8300; 10 nm excitation and emission bandwidth; 8 s integration time) around the emission maxima of the fluorophores (for donor, λ_ex_ = 535 nm and λ_em_ = 580 nm; for acceptor, λ_ex_ = 635 nm and λ_em_ = 660 nm). Anisotropy values rwere obtained from on r= (*l*_*VV*_ -*Gl*_*H*_)/(*l*_*VV*_ + 2*Gl*_*H*_), where *l*_*VV*_and *l*_*H*_ are the fluorescence emission intensities in the vertical and horizontal orientation, respectively, upon excitation along the vertical orientation. The sensitivity of the spectrometer to different polarizations was corrected via the factor *G=l*_*H*_*l*_*HH*_, where *l*_*H*_and *l*_*HH*_are the fluorescence emission intensities in the vertical and horizontal orientation, respectively, upon excitation along the horizontal orientation. *G*-values were determined to be 1.8-1.9. The anisotropy was measured in buffer A and the labelled proteins and free-fluorophores in a concentration range of 50-500 nM at room temperature.

### Solution-based smFRET and ALEX

Solution-based smFRET and alternating laser excitation (ALEX)^49^ experiments were carried out at 25-100 pM of labelled protein at room temperature in the appropriate buffer (50 mM Tris-HCl, pH 7.4; 50 mM KCl for MalE and OppA; 25 mM Tris-HCl, pH 8.0; 150 mM NaCl; 1 μM EDTA for PsaA; 50 mM KPi, pH 7.4; 50 mM KCl for OpuAC, SBD1 and SBD2) supplemented with additional reagents as stated in the text. Microscope cover slides (no. 1.5H precision cover slides, VWR Marienfeld) were coated with 1 mg/mL BSA for 30-60 s to prevent fluorophore and/or protein interactions with the glass material. Excess BSA was subsequently removed by washing and exchange with appropriate buffer (50 mM Tris-HCl, pH 7.4; 50 mM KCl for MalE and OppA; 25 mM Tris-HCl, pH 8.0; 150 mM NaCl; 1 μM EDTA for PsaA; 50 mM KPi, pH 7.4; 50 mM KCl for OpuAC, SBD1, SBD2).

All smFRET experiments were performed using a home-built confocal microscope. In brief, two laser-diodes (Coherent Obis) with emission wavelength of 532 and 637 nm were directly modulated for alternating periods of 50 µs and used for confocal excitation. The laser beams where coupled into a single-mode fiber (PM-S405-XP, Thorlabs) and collimated (MB06, Q-Optics/Linos) before entering a water immersion objective (60X, NA 1.2, UPlanSAPO 60XO, Olympus). The fluorescence was collected by excitation at a depth of 20 µm. Average laser powers were 30 μW at 532 nm (∼30 kW/cm^2^) and 15 μW at 637 nm (∼15 kW/cm^2^). Excitation and emission light was separated by a dichroic beam splitter (zt532/642rpc, AHF Analysentechnik), which is mounted in an inverse microscope body (IX71, Olympus). Emitted light was focused onto a 50 µm pinhole and spectrally separated (640DCXR, AHF Analysentechnik) onto two single-photon avalanche diodes (TAU-SPADs-100, Picoquant) with appropriate spectral filtering (donor channel: HC582/75; acceptor channel: Edge Basic 647LP; AHF Analysentechnik). Photon arrival times in each detection channel were registered by an NI-Card (PXI-6602, National Instruments) and processed using custom software implemented in LabView (National Instruments).

An individual labelled protein diffusing through the confocal volume generates a burst of photons. To identify fluorescence bursts a dual-colour burst search^67^ was used with parameters M = 15, T = 500 μs and L = 25. In brief, a fluorescent signal is considered a burst, when a total of L photons having M neighbouring photons within a time window of length T centred on their own arrival time. A first burst search was done that includes the donor and acceptor photons detected during the donor excitation, and a second burst search was done including only the acceptor photons detected during the acceptor excitation. The two separate burst searches were combined to define intervals when both donor and acceptor fluorophores are active. These intervals define the bursts. Only bursts having >150 photons were further analysed

The three relevant photon streams were analysed (DA, donor-based acceptor emission; DD, donor-based donor emission; AA, acceptor-based acceptor emission) and assignment is based on the excitation period and detection channel^49^. The apparent FRET efficiency is calculated via F(DA)/[F(DA)+F(DD)] and the Stoichiometry S by [F(DD)+F(DA)]/[(F(DD)+F(DA)+F(AA)], where F(·) denotes the summing over all photons within the burst^49^.

Binning the detected bursts into a 2D apparent FRET/S histogram (81 × 81 bins) allowed the selection of the donor and acceptor labelled molecules and reduce artefacts arising from fluorophore bleaching^49^. The selected apparent FRET histogram were fitted with a Gaussian distribution using nonlinear least square, to obtain a 95% Wald confidence interval for the distribution mean. Significance statements about the mean of the FRET distributions were made by comparing appropriate confidence intervals.

### Scanning confocal microscopy

Confocal scanning experiments were performed at room temperature and using a home-built confocal scanning microscope as described previously^29, 68, 69^. In brief, surface scanning was performed using a XYZ-piezo stage with 100×100×20 µm range (P-517-3CD with E-725.3CDA, Physik Instrumente). The detector signal was registered using a Hydra Harp 400 picosecond event timer and a module for time-correlated single photon counting (both Picoquant). Data were recorded with constant 532 nm excitation at an intensity of 0.5 μW (∼125 W/cm^2^) for SBD2, PsaA, OppA and MalE, but 1.5 μW (∼400 W/cm^2^) for OpuAC, unless stated otherwise. Scanning images of 10×10 µm were recorded with 50 nm step size and 2 ms integration time at each pixel. After each surface scan, the positions of labelled proteins were identified manually; the position information was used to subsequently generate time traces. Surface immobilization was conducted using an anti-HIS antibody and established surface-chemistry protocols as described^29^. A flow-cell arrangement was used as described before^29, 70^ for studies of surface-tethered proteins, except for MalE. MalE was studied on standard functionalized cover-slides since MalE was extremely sensitive to contaminations of maltodextrins in double-sided tape or other flow-cell parts. All experiments of OpuAC and PsaA were carried out in degassed buffers (50 mM KPi pH 7.4, 50 mM KCl for OpuAC and 25 mM Tris-HCl pH 8.0, 150 mM NaCl, 1 μM EDTA for PsaA) under oxygen-free conditions obtained utilizing an oxygen-scavenging system supplemented with 10 mM of Trolox (Merck)^71^. For MalE, SBD1, SBD2 and OppA experiments were carried out in buffer (50 mM KPi, pH 7.4, 50 mM KCl for SBD2 and 50 mM Tris-HCl, pH 7.4, 50 mM KCl for MalE and OppA) supplemented with 1 mM Trolox and 10 mM Cysteamine (Merck).

### Analysis of fluorescence trajectories

Time-traces were analysed by integrating the detected red and green photon streams in time-bins as stated throughout the text. Only traces lasting longer than 50 time-bins, having on average more than 10 photons per time-bin that showed clear bleaching steps, were used for further analysis. The number of analysed molecules, transitions and the total observation time are indicated in Table 4. The apparent FRET per time-bin was calculated by dividing the red photons by the total number of photons per time-bin. The state-trajectory of the FRET time-trace was modelled by a Hidden Markov Model (HMM)^72^. For this an implementation of HMM was programmed in Matlab (MathWorks), based on the work of Rabiner^72^. In the analysis, we assumed that the FRET time-trace (the observation sequence) can be considered as a HMM with two states having a one-dimensional Gaussian-output distribution. The Gaussian output-distribution of state *i*(*i* =1, 2) was completely defined by the mean and the variance. The goal was to find the parameters λ (transition probabilities that connect the states and parameters of output-distribution), given only the observation sequence that maximizes the likelihood function. This was iteratively done using the Baum-Welch algorithm^73^. Care was taken to avoid floating point underflow and was done as described^72^. With the inferred parameters λ, the most probable state-trajectory is then found using the Viterbi algorithm^74^. The time spent in each state (open, closed) was inferred from the most probable state-trajectory, an histogram was made and the mean time spent in each state was calculated.

### Ensemble FRET

Fluorescence spectra of labelled SBD1 and SBD2 proteins were measured on a scanning spectrofluorometer (Jasco FP-8300; λ_ex_ = 552 nm, 5 nm excitation and emission bandwidth; 3 s integration time). The apparent FRET efficiency was calculated via I_Acceptor_/(I_Acceptor_+ I_donor_), where I_Acceptor_ and I_donor_ are fluorescence intensities around the emission maxima of the acceptor (660 nm) and donor fluorophore (600 nm), respectively. Measurements were performed at 20°C with ∼200 nM labelled protein dissolved in buffer A.

## ACKNOWLEGDEMENTS

This work was financed by an NWO Veni grant (722.012.012 to G.G.), an ERC Advanced Grant (No. 670578 – ABCvolume to B.P.), the National Health and Medical Research Council (Project Grants 1080784 and 1122582 to C.A.M), the Australian Research Council (Discovery Project DP170102102 and Future Fellowship FT170100006 to C.A.M.) and an ERC Starting Grant (No. 638536 – SM-IMPORT to T.C.). G.G. also acknowledges an EMBO fellowship (long-term fellowship ALF 47-2012 to G.G.) and financial support by the Zernike Institute for Advanced Materials. G.G. is a Rega foundation post-Doctoral fellow. T.C. was further supported by the Center of Nanoscience Munich (CeNS), Deutsche Forschungsgemeinschaft within GRK2062/1 (project C03) and SFB863 (project A13), LMUexcellent and the Center for integrated protein science Munich (CiPSM). We thank H. Jung, D. Griffith and M. Wiertsema for reading of the manuscript.

## AUTHOR CONTRIBUTIONS

M.d.B., G.G., B.P., C.A.M. and T.C. designed the study. B.P., C.A.M. and T.C. supervised the project. M.d.B., G.G., R.V. and F.H. performed the molecular biology and protein chemistry studies and developed the labeling protocols. M.d.B., G.G., R.V., F.H., and N.E. performed single-molecule experiments. G.K.S. performed transport assays and ITC. M.d.B. analysed smFRET data. S.L.B. and C.A.M. designed and executed the PsaA biochemical studies. All authors contributed to discussion of the research and writing of the manuscript.

## AUTHOR INFORMATION

The authors declare no competing financial interest. Correspondence and requests for material should be addressed to B.P., C.A.M. and T.C.

---

**Figure 2S1.**
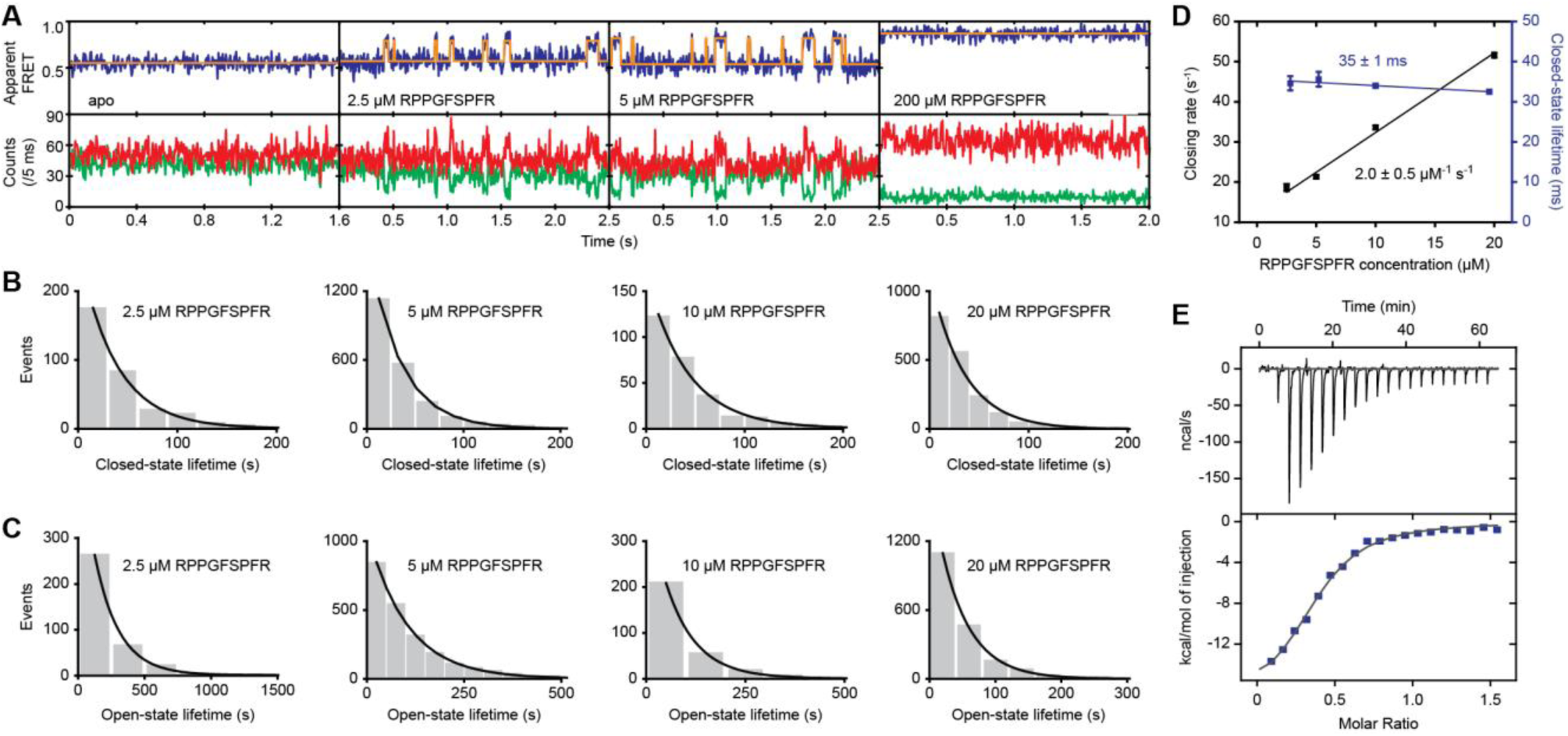
figure supplements 1. OppA uses an induced-fit ligand binding mechanism. (**A**) Representative fluorescence trajectories of OppA(A209C/S441C) at different peptide (RPPGFSFR) concentrations; donor (green) and acceptor (red) photon counts. The top panel shows the calculated apparent FRET efficiency (blue) with the most probable state-trajectory of the Hidden Markov Model (HMM) (orange). Dwell time histogram of the low FRET (closed conformation) (**B**) and high FRET state (open conformation) (**C**) as obtained from the most probable state-trajectory of the HMM of all molecules per condition. Grey bars are the binned data and the solid line is an exponential fit. Total number of analysed molecules are indicated in Table 4. (**D**) Closing rate (rate of low to high FRET state; black) and lifetime of the ligand-bound conformation (lifetime high FRET state; purple) of OppA as obtained from the most probable state-trajectory of the HMM of all molecules at different peptide (RPPGFSFR) concentrations. Data correspond to mean ± s.e.m. and the solid line a linear fit. Slope or intercept of the fit are shown (95% confidence interval). From the fit a dissociation constant K_D_ of 14 ± 4 µM (95% confidence interval) is obtained. (**E**) Isothermal calorimetry binding isotherms of the titration of OppA with the peptide (RPPGFSFR), which yielded a dissociation constant K_D_ of 5 ± 3 µM (mean ± s.d., n = 4). Points are the data and the solid line a fit to a one site-binding model.

**Figure 2S2.**
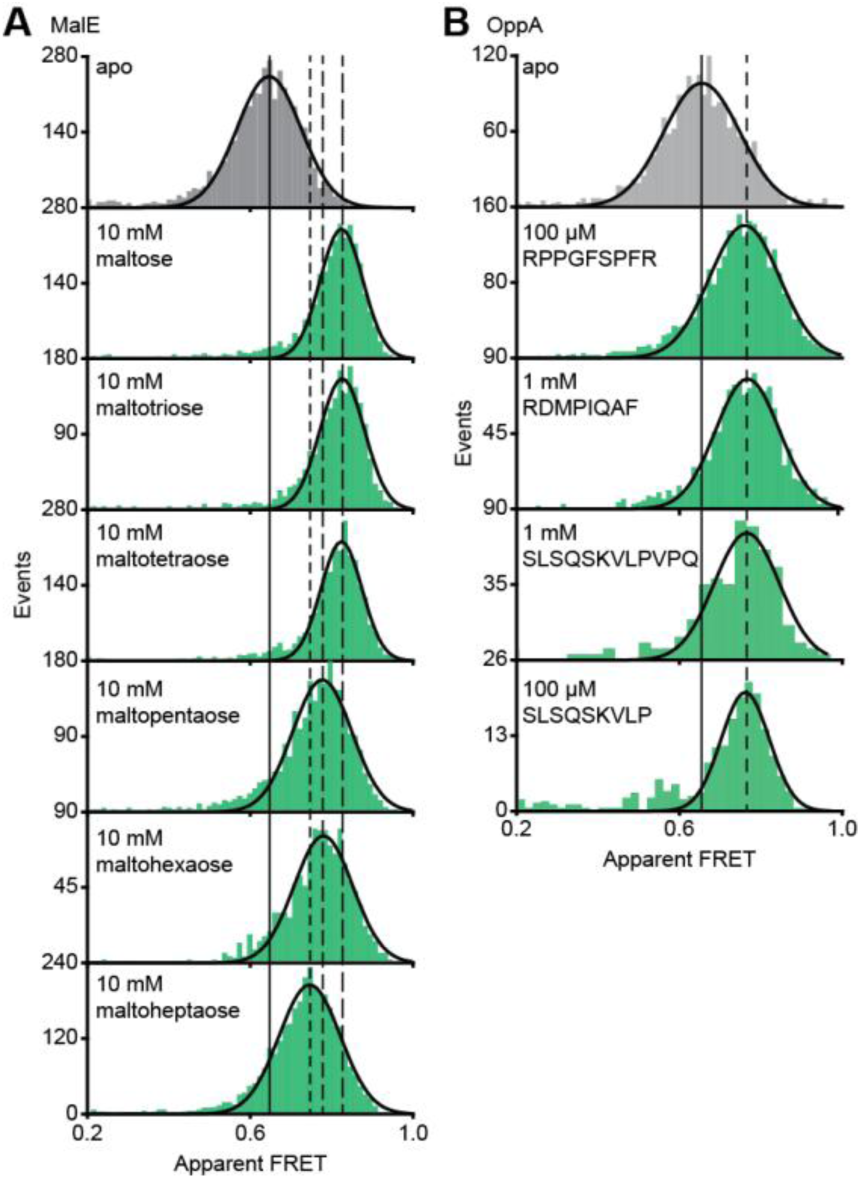
figure supplements 2. Translocation competent conformation(s) of MalE and OppA. Solution-based apparent FRET efficiency histogram of MalE(T36C/S352C) (**A**) and OppA(A209C/S441C) (**B**) in the absence and presence of different cognate substrates as indicated. The OppA substrates are indicated by one-letter amino acid code. Bars are experimental data and the solid line a Gaussian distribution fit. The 95% confidence interval for the mean of the Gaussian distribution is shown in Table 3, and the interval centre is indicated by vertical lines (solid and dashed).

**Figure 2S3.**
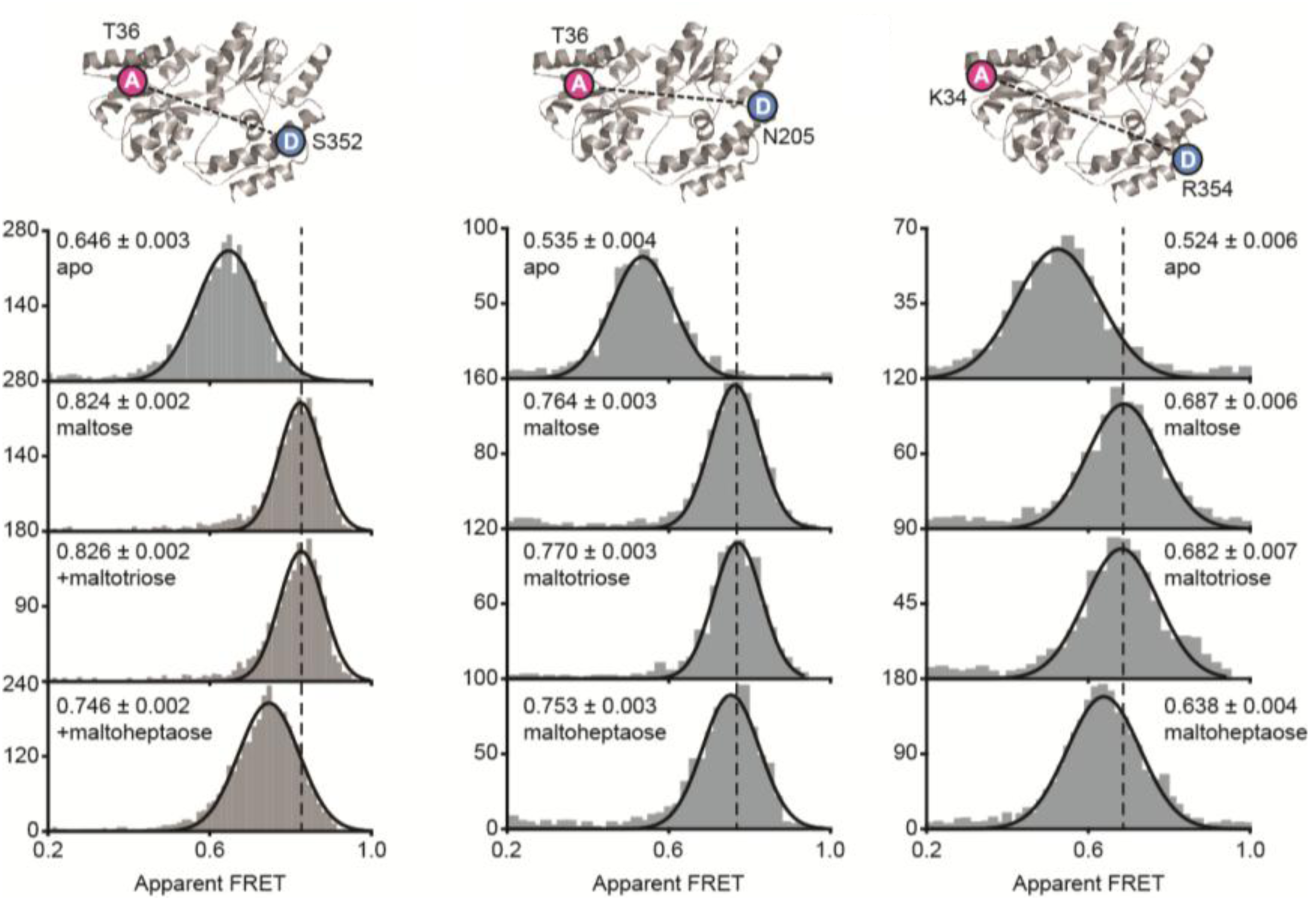
figure supplements 3. MalE conformations studied by smFRET. Solution-based apparent FRET efficiency histogram of MalE(T36C/S352C), MalE(T36C/N205C) and MalE(K34C/R354C) in the absence and presence of different cognate substrates as indicated. Bars are experimental data and the solid line a Gaussian distribution fit. The 95% confidence interval for the mean of the Gaussian distribution is shown in Table 3, and the interval centre is indicated by vertical lines (solid and dashed). Structure of ligand-free MalE (PDB ID: 1OMP) with corresponding donor and acceptor fluorophore positions is indicated above the histograms.

**Figure 3S1.**
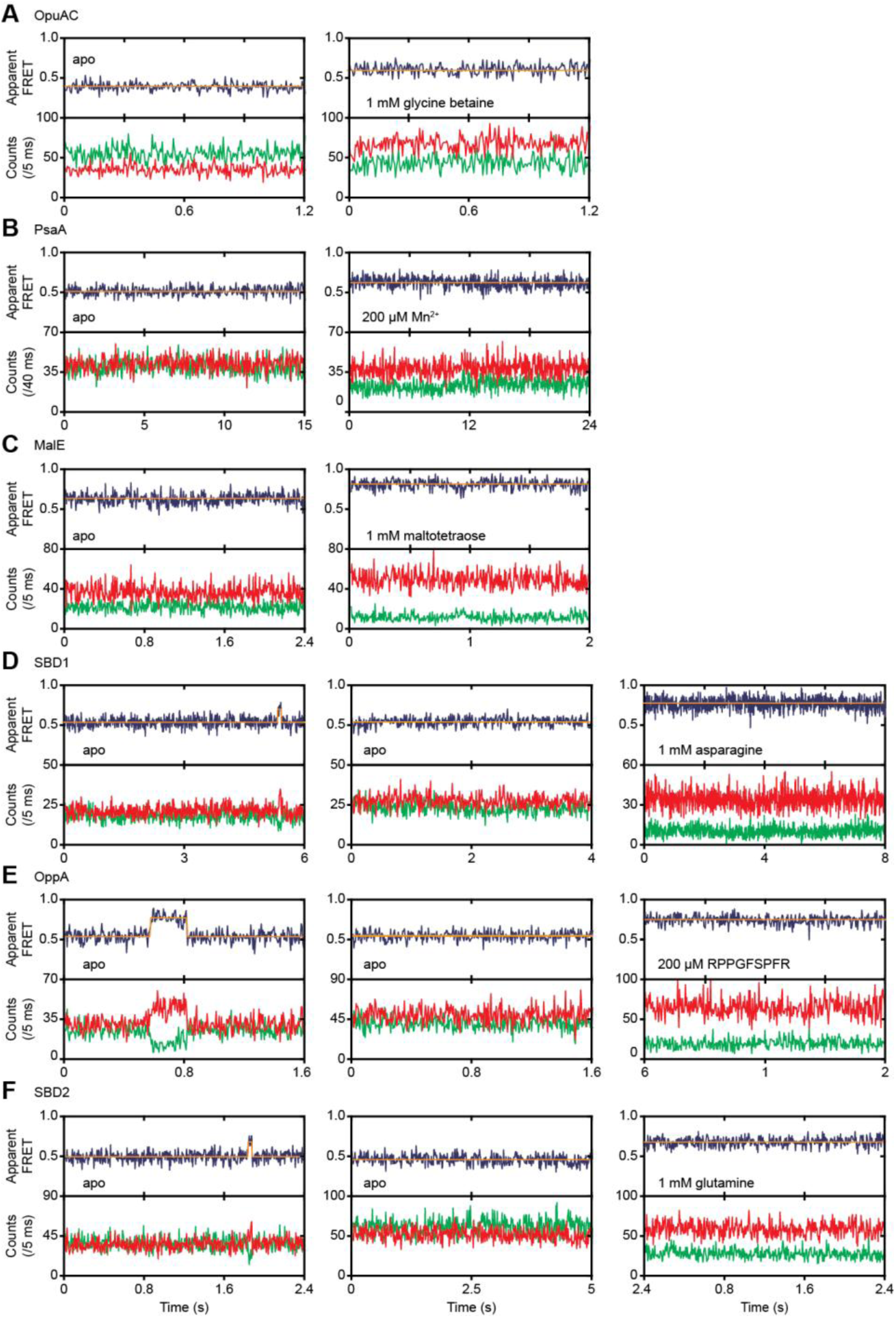
figure supplements 1. Conformational dynamics of ligand-free and ligand-bound SBPs. Representative fluorescence trajectories of OpuAC(V360C/N423C) (**A**), PsaA(V76C/K237C) (**B**), MalE(T36C/S352C) (**C**), SBD1(T159C/G87C) (**D**), OppA(A209C/S441C) (**E**) and SBD2(T369C/S451) (**F**) in the absence of substrate and under saturating conditions of ligand, as indicated. In the absence of ligand, 20 μM of unlabelled protein or 1 mM EDTA (for PsaA) was added to scavenge any ligand contaminations. The top panels show the calculated apparent FRET efficiency (blue) from the donor (green) and acceptor (red) photon counts as presented in bottom panels. Orange line indicate average apparent FRET efficiency value or most probable state-trajectory of the HMM. Statistics can be found in Table 4.

**Figure 4S1.**
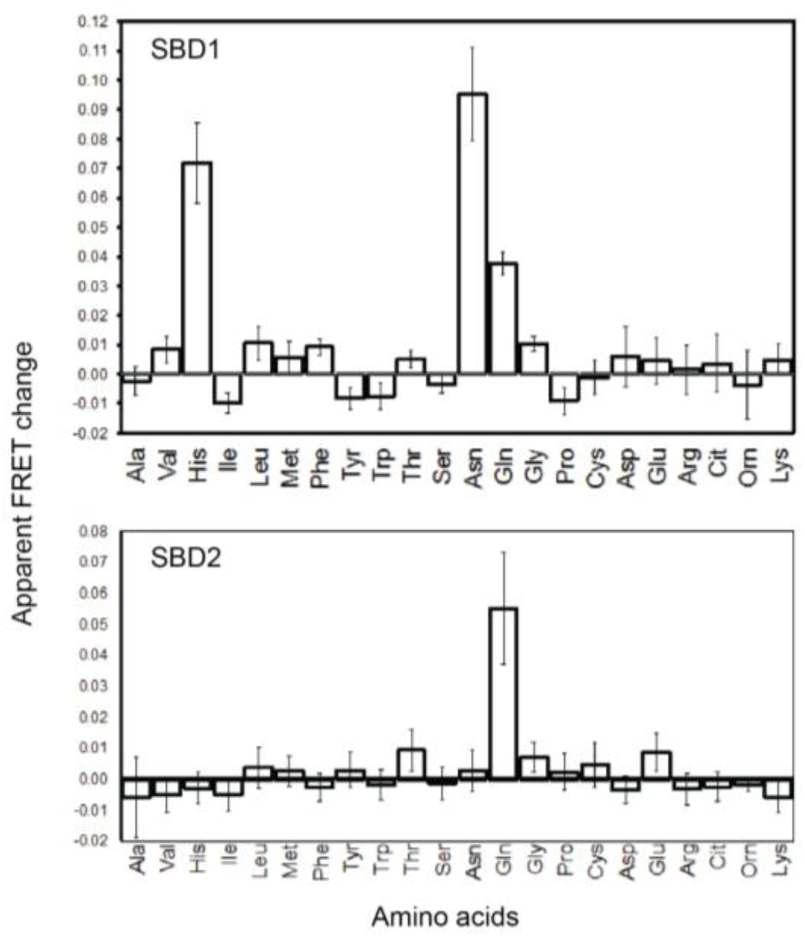
figure supplements 1. Substrate binding of SBD1 and SBD2 studied by ensemble FRET. The mean apparent FRET change of SBD1 (top) and SBD2 (bottom) in the presence of 5 mM of the indicated amino acids relative to their absence; measurements were performed in 50 mM KPi, 50 mM KCl, pH 7.4. Amino acids are indicated by their three letter abbreviation. Data correspond to mean ± s.d. of the apparent FRET change of duplicate measurements with the same labelled protein sample.

**Figure 4S2.**
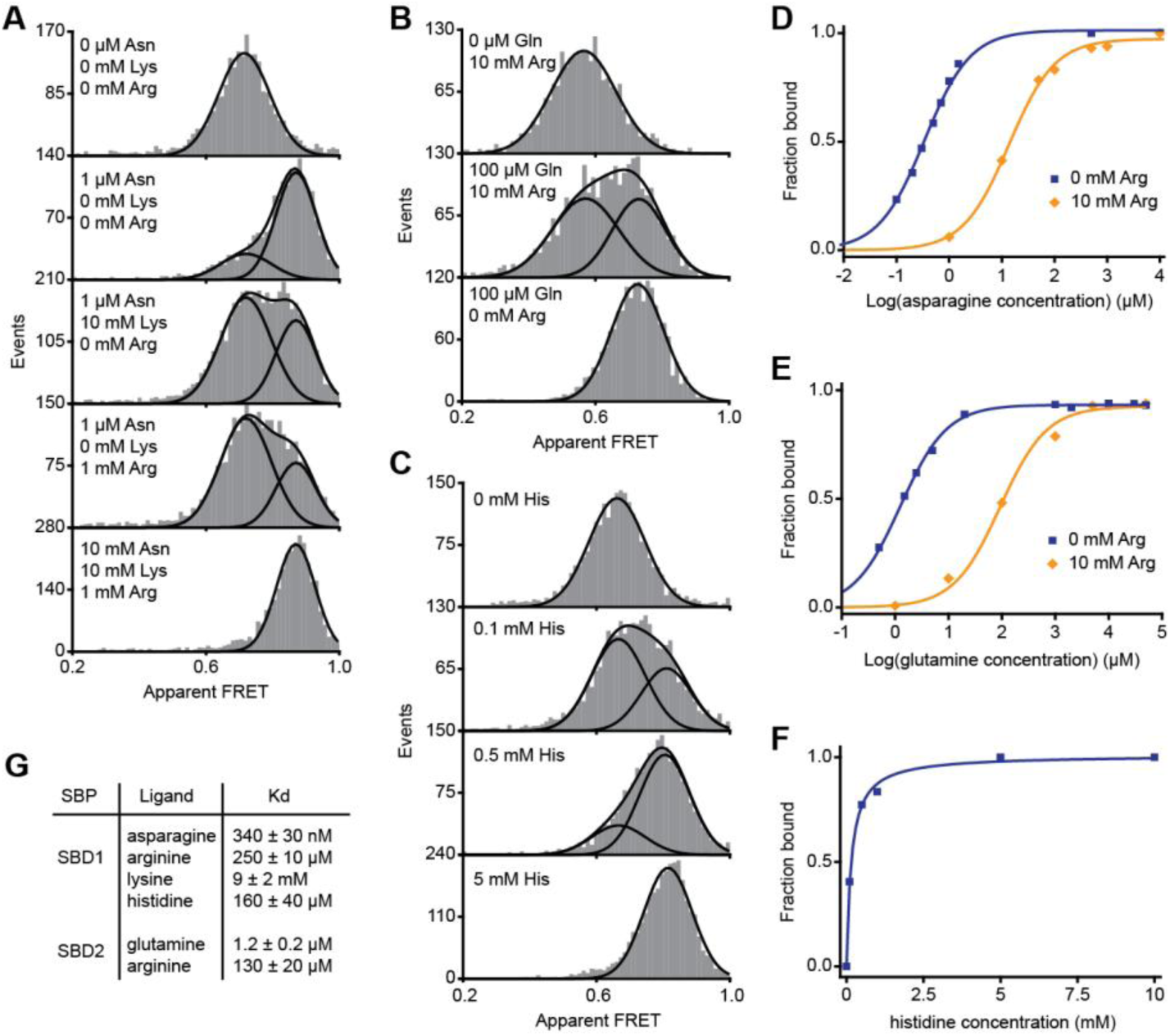
figure supplements 2. Non-cognate substrate binding by SBD1 and SBD2. Solution-based apparent FRET efficiency histograms of SBD1(T159C/G87C) (**A** and **C**) and SBD2(T369C/S451) (**B**) in the presence of different ligand concentrations as indicated. Bars are experimental data and the solid lines a fit to a mixture model with two Gaussian distributions or a fit with a single Gaussian distribution. The mean of the Gaussian distributions was obtained from the extreme conditions and fixed in the mixture model. Fraction of SBD1 bound to asparagine (**D**), SBD2 bound to glutamine (**E**) and SBD1 bound to histidine (**F**). Points are the data and the solid line a fit to a one site-binding model. (**G**) Estimated dissociation constants K_D_ as obtained from the fit. Error bars represent 95% confidence interval.

**Figure 6S1.**
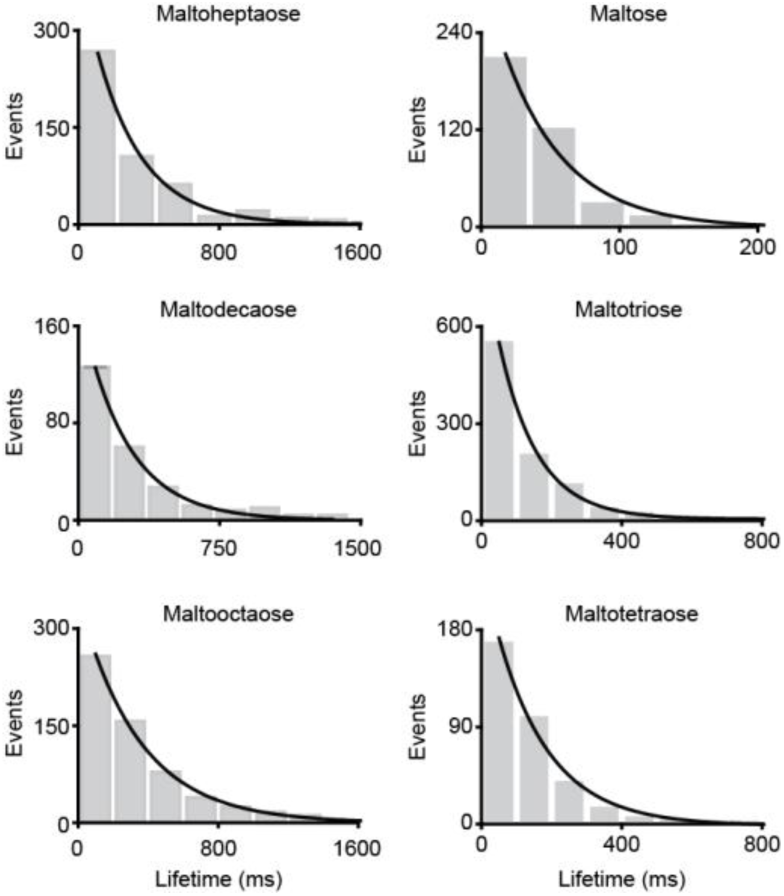
figure supplements 1. Distribution of the ligand-bound conformations of MalE. Dwell time histogram of the high FRET (closed ligand-bound conformation) as obtained from the most probable state-trajectory of the HMM of all molecules per condition as shown in Figure 6. Grey bars are the binned data and the solid line an exponential fit. Statistics can be found in Table 4.

**Figure 6S2.**
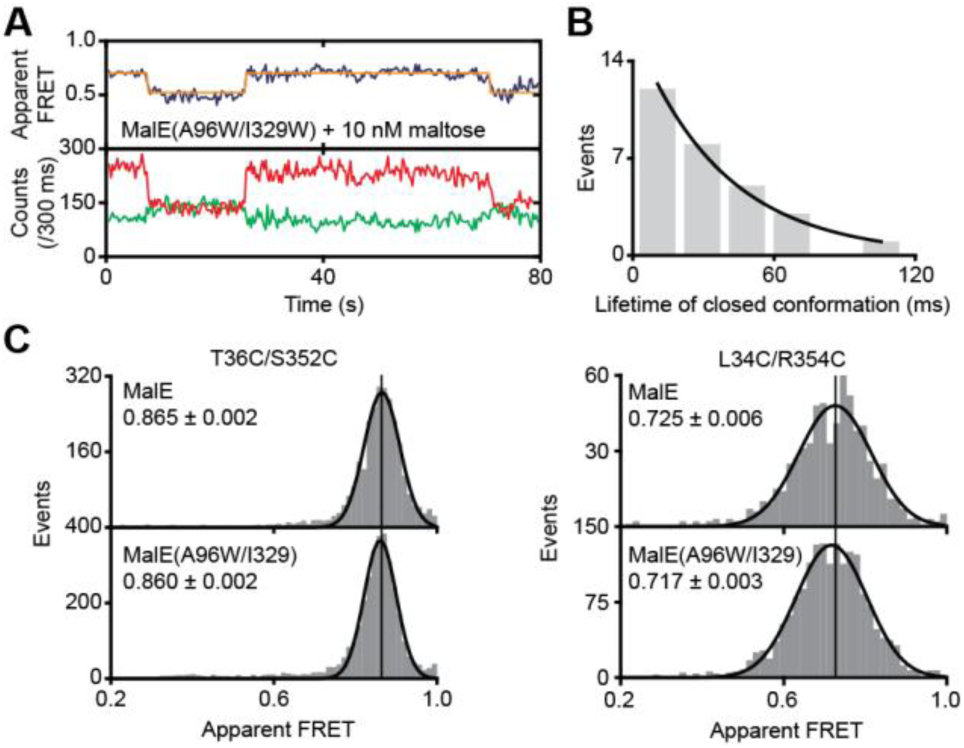
figure supplements 2. Conformational changes and dynamics of MalE(A96W/I329W). (**A**) Representative fluorescence trajectories of MalE(T36C/S352C/A96W/I329W) in the presence of 10 nM maltose. Fluorescence trajectories: the top panel shows the calculated apparent FRET efficiency (blue) from the donor (green) and acceptor (red) photon counts as shown in the bottom panel. The most probable state-trajectory of the Hidden Markov Model (HMM) is shown (orange). (**B**) Dwell time histogram of the high FRET state (closed conformation) as obtained from the most probable state-trajectory of the HMM of all molecules. Grey bars are the binned data and the solid line is an exponential fit. Statistics can be found in Table 4. (**C**) Solution-based apparent FRET efficiency histogram of MalE and MalE(A96W/I32W) in the presence of 1 mM maltose for the indicated inter-dye positions. Bars are experimental data and solid line a Gaussian distribution fit. The 95% confidence interval for the mean of the Gaussian distribution is indicated.

